# A method for independent estimation of false localisation rate for phosphoproteomics

**DOI:** 10.1101/2021.10.18.464791

**Authors:** Kerry A Ramsbottom, Ananth Prakash, Yasset Perez Riverol, Oscar Martin Camacho, Maria Martin, Juan Antonio Vizcaíno, Eric W Deutsch, Andrew R Jones

## Abstract

Phosphoproteomics methods are commonly employed in labs to identify and quantify the sites of phosphorylation on proteins. In recent years, various software tools have been developed, incorporating scores or statistics related to whether a given phosphosite has been correctly identified, or to estimate the global false localisation rate (FLR) within a given data set for all sites reported. These scores have generally been calibrated using synthetic data sets, and their statistical reliability on real datasets is largely unknown. As a result, there is considerable problem in the field of reporting incorrectly localised phosphosites, due to inadequate statistical control.

In this work, we develop the concept of using scoring and ranking modifications on a decoy amino acid, i.e. one that cannot be modified, to allow for independent estimation of global FLR. We test a variety of different amino acids to act as the decoy, on both synthetic and real data sets, demonstrating that the amino acid selection can make a substantial difference to the estimated global FLR. We conclude that while several different amino acids might be appropriate, the most reliable FLR results were achieved using alanine and leucine as decoys, although we have a preference for alanine due to the risk of potential confusion between leucine and isoleucine amino acids. We propose that the phosphoproteomics field should adopt the use of a decoy amino acid, so that there is better control of false reporting in the literature, and in public databases that re-distribute the data. Data are available via ProteomeXchange with identifier PXD028840.

## Introduction

There is great research interest in studying post-translational modifications (PTMs) to proteins, due to their importance in cell signalling, as a rapid mode of proteins changing their function, and their implication in almost all known disease processes. The most widely studied reversible modifications include phosphorylation (by far the most studied one, and our primary focus here), acetylation, methylation, and attachment of small proteins, such as ubiquitin and SUMO.

High throughput tandem mass spectrometry (MS) is commonly used for detection and localisation of phosphorylation sites on proteins, using so-called phosphoproteomics methods. Typically in these methods, proteins are first extracted from samples, digested with an enzyme such as trypsin and then phosphorylated peptides are enriched in a sample, for example using TiO2 or other metal ion attached to a column (affinity chromatography), to which phosphate binds preferentially. Bound peptides are then eluted and analysed by liquid chromatography-mass spectrometry (LC-MS/MS) [1]. In the common analysis mode used in phosphoproteomics, data dependent acquisition (DDA) is performed to fragment the most abundant peptides observed. The MS^2^ fragmentation spectra (plus the mass/charge of the intact precursor) are then used to identify peptide sequences, for example using sequence database search software. In this approach, the spectra are searched against a theoretical digest of the proteome (i.e. peptide sequence database) for the given species, taking account for the variable modifications selected. For phosphorylation, most users search for phosphorylation on the canonical Ser, Thr and Tyr (STY) residues, where the vast majority of detectable phosphorylation resides in eukaryotic systems. The search engine then considers every STY residue with and without the addition of the phosphate mass (+79.97 Daltons), greatly increasing the size of the search space, with a corresponding reduction in statistical power for peptide identification. Confident peptide identification is governed by the quality of the match between the observed spectrum and the theoretical spectrum expected for a peptide from the sequence database, from which local statistics such as p-values or e-values are usually calculated, as well as sometimes a PEP (posterior error probability). If a PEP value is calculated, 1-PEP gives the probability that a given peptide-spectrum match (PSM) is correct. There are of course many different proteomics search engines, including commercial and free and/or open source, for a review see Verheggen *et al*. [2].

An important consideration for phosphoproteomics is the confidence that a given site within a protein has been correctly identified as being phosphorylated. Ambiguity in this regard may occur when a confident PSM has more STY residues than *n*, where *n* is the number of phosphorylation modification instances detected i.e. intact peptidoform mass = peptide sequence mass + (n * 79.97 Da). In this case, the search engine itself, or a downstream analysis package, calculates statistics related to each of the *n* phosphosites within a peptide, such as a PEP that the site has been incorrectly localised (sometimes also called local false localisation rate; local FLR) or other *ad hoc* score. As for PSMs, if an accurate PEP can be estimated, then 1-PEP_site_ gives the probability that the site has been correctly localised, in this case assuming already that the PSM is definitively correct. Correct site localisation can be critically important for downstream uses of data. As one example, there are completely different kinases and phosphatases involved in Ser/Thr versus Tyr phosphorylation, and thus biological conclusions as to the up and downstream signalling pathways would be completely different. Even where ambiguity relates to different, say, Ser residues in a peptide, nearby amino acid motifs allow inference of the kinase responsible for phosphorylating the site, and thus incorrect site determination could lead researchers into making incorrect assumptions and conclusions.

Many of the published site localisation algorithms were benchmarked by the originating authors and scores calibrated based on synthetic data sets, with a known “ground truth” i.e. where the sites of phosphorylation were known [3]. There have also been some independent efforts to benchmark different site localisation tools, showing that the choice of tool does alter the global statistics [4, 5] i.e. sensitivity - how many sites in a whole data set can be correctly localised at a suitable overall (global) FLR. While tools continue to improve and become more widely used for ensuring confident site localisation, there remain several unsolved challenges for the field, as follows. First, it is unclear whether findings on synthetic data sets can be extrapolated to genuine biological data sets, with generally a higher level of complexity. Second, for analysis of real experimental data sets, there are no commonly used methods for independent estimation of *global FLR*, for example that allow a researcher to ask the question – how many sites have we confidently identified and how many are likely false positives in the whole data set. For regular peptide/protein identification in proteomics, decoy database search methods are now almost ubiquitously used for estimating peptide/protein false discovery rate (FDR), since they give a search engine independent statistic that is easy to understand. There is not a generally accepted method for calculating the same type of statistic for PTM site identification. Third, our groups are interested in very large-scale re-analysis of public PTM enriched data sets, via a project called PTMeXchange. We wish to have methods that allow for accurate calculation of the probability that a given PTM site has been observed in a meta-analysis of data sets, where there could be potentially multiple PSMs from different studies supporting a given site. To our knowledge, there are no suitable statistical models for combining different evidence streams.

In this work, we explore the concept of using decoy amino acid(s) for estimation of site localisation statistics (global FLR), in this context defined as one that we know cannot be modified, to model the distribution of false localisations detected from a processing pipeline. We test a range of different amino acids for their suitability as a decoy in synthetic and real data sets, as well as demonstrating the results obtained from several common proteome informatics pipelines. The concept of using a decoy amino acid for localisation of PTMs is not a new one. It has been used before in several previous publications and approaches [6, 7]. However, to our knowledge, no publication has yet validated the statistics associated with the use of decoy amino acids, particularly on multiple tools, and the method of using a decoy amino acid has not gained widespread use in the field. The majority of PTM-based studies still rely on using *ad hoc* score thresholds for determining whether PTMs have been correctly identified or not. From the results we present, we make some recommendations as to how we believe large-scale PTM-enriched studies should be analysed to control the local and global false localisation rate. While we have focussed on computational analysis and pipelines for phosphorylation, general approaches and conclusions are largely applicable to other types of PTM readily detected by mass spectrometry (MS). Code used for the analysis is in GitHub: https://github.com/PGB-LIV/PhosphoFLR.

## Methods

Our overall goal is to demonstrate methods for controlling and understanding FLR, rather than benchmarking tools *per se*, although we wished to demonstrate the reproducibility of methods in different pipelines. As such, we tested four commonly used analysis pipelines: Trans-Proteomic Pipeline (TPP) [8] including Comet search [9] and PTMProphet site localization [10]; MaxQuant including PTMScore [11]; ProteomeDiscover, including Mascot search and ptmRS localization [12, 13]; and PEAKS DB search with A-Score [14]. We tested the effects on global FLR of selecting different amino acids as the “decoy”, and profiled the frequency of potential decoy amino acids relative to assumed correct STY phosphorylation sites, to see which provides the decoy distribution best matching the target distribution i.e. other STY sites to which the site could be wrongly localized. The mass spectrometry proteomics data have been deposited to the ProteomeXchange Consortium via the PRIDE [15] partner repository with the dataset identifier PXD028840 and DOI 10.6019/PXD028840

Three data sets were used for evaluation of methods for estimating global FLR – one synthetic data set, one model plant phosphoproteomics data set (from *Arabidopsis thaliana*) and one human phosphoproteomics data set. The raw files of the three search data sets were obtained from the ProteomeXchange Consortium [16] via the PRIDE repository [17]. These included ten files from the PXD007058 [5] synthetic data set (files named “HCDOT” pools 1 to 5, reps 1 and 2), twelve files from the PXD008355 [18] *Arabidopsis* data set (rapamycin treated) and six from the PXD000612 [19] human data set (files randomly selected). The PXD007058 data set contains a synthetic phosphopeptide library. The use of synthetic phosphopeptides allowed us to define FLR (through one method) by comparing the results from our search pipelines to the known phosphopeptide sequences to determine if our analyses correctly localise the phosphosites. The PXD008355 *Arabidopsis thaliana* data set and the PXD000612 human data set are both biological data sets with unknown phosphosites.

Databases were created for the searches of each data set. The PXD007058 search database consisted of the synthetic peptides [5], the PXD008355 *Arabidopsis* search database of Araport11 [20] sequences and the PXD000612 human search database was created from the Level 1 PeptideAtlas Tiered Human Integrated Search Proteome [21], containing core isoforms from neXtProt [22]. Each search database also contained the cRAP contaminants sequences (https://www.thegpm.org/crap/, last accessed October 2021). Decoys across all three databases were generated for each entry using the Brujin method (with k=2) [23].

Using the Trans-Proteomic Pipeline (TPP) [8], the data set files were first searched using Comet [9]. The resulting files were then combined and processed using PeptideProphet [24], iProphet [25], and PTMProphet [10]. In addition to searching for phosphorylation, we also searched for pyrophosphorylation modifications which pilot searches had determined were present on some synthetic peptides. The key Comet search parameters for each dataset are shown in Table 1; the key parameters used for the other pipelines are shown in Supplementary Table 1.

**Table 1:**
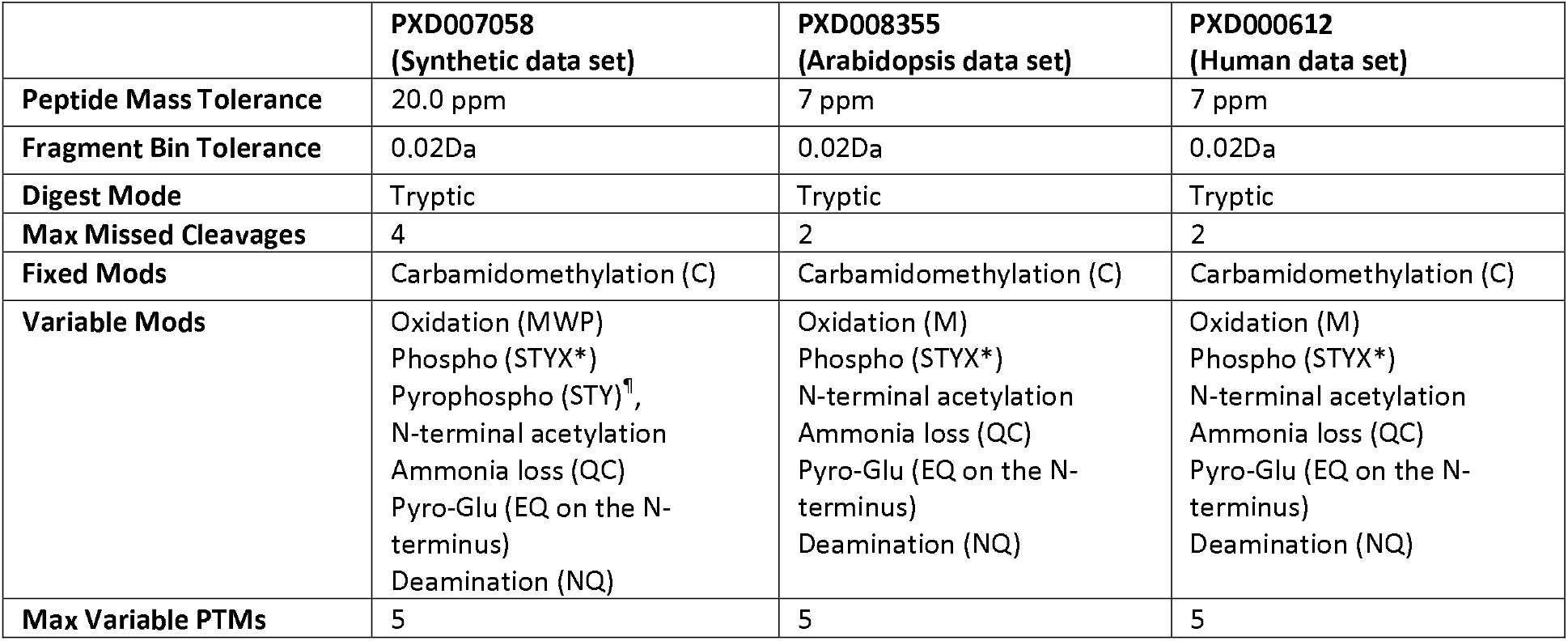
Comet search params for each data set. *X corresponds to the different decoy amino acid searched: Ala, Gly, Leu, Asp, Glu or Pro.^¶^ Preliminary analysis of the data set detected that several peptides had been manufactured with a pyrophosphate modification rather than the intended phosphate, which can cause apparent errors when comparing the results to the answer key if they are not accounted for.

|  | PXD007058<br>(Synthetic data set) | PXD008355<br>(Arabidopsis data set) | PXD000612<br>(Human data set) |
| --- | --- | --- | --- |
| **Peptide Mass Tolerance** | 20.0 ppm | 7 ppm | 7 ppm |
| **Fragment Bin Tolerance** | 0.02Da | 0.02Da | 0.02Da |
| **Digest Mode** | Tryptic | Tryptic | Tryptic |
| **Max Missed Cleavages** | 4 | 2 | 2 |
| **Fixed Mods** | Carbamidomethylation (C) | Carbamidomethylation (C) | Carbamidomethylation (C) |
| **Variable Mods** | Oxidation (MWP)<br>Phospho (STYX*)<br>Pyrophospho (STY)<sup>¶</sup>,<br>N-terminal acetylation<br>Ammonia loss (QC)<br>Pyro-Glu (EQ on the N-terminus)<br>Deamination (NQ) | Oxidation (M)<br>Phospho (STYX*)<br>N-terminal acetylation<br>Ammonia loss (QC)<br>Pyro-Glu (EQ on the N-terminus)<br>Deamination (NQ) | Oxidation (M)<br>Phospho (STYX*)<br>N-terminal acetylation<br>Ammonia loss (QC)<br>Pyro-Glu (EQ on the N-terminus)<br>Deamination (NQ) |
| **Max Variable PTMs** | 5 | 5 | 5 |

### Downstream data analysis

The data from searching with TPP were downstream processed by custom Python scripts (https://github.com/PGB-LIV/PhosphoFLR). Firstly, the global FDR was calculated from the decoy counts and the PSMs were filtered for 1% PSM FDR. From these filtered PSMs, a site-based file was generated giving separate localisation scores for each phosphosite found on each PSM, removing decoy and contaminant hits. These site-based PSMs were ordered by a *combined probability*, calculated by multiplying the PSM probability by the localisation probability. In the processing pipeline from TPP, iProphet calculates a probability that a given PSM is correct, and PTMProphet calculates a probability for the site assignment. We demonstrated that there is almost no meaningful correlation (r^2^ =~ 0.01) between these probabilities (Supplementary Figure 1), and thus we conclude that these probabilities are sufficiently independent that they can be multiplied to arrive at a final probability that a given site’s identification is supported by the given spectrum.

For the PEAKS search, the PSM score and A-scores for targets and decoys were modelled based on the counts of targets and decoys per histogram score bin, to generate similar probability estimates (code provided in the GitHub repository). For MaxQuant and Mascot searches, the PSM probability values were calculated as 1-PEP values (reported by the pipeline natively) with the PTM probabilities being calculated innately through PTM-score/ptmRS probabilities, respectively. For the synthetic peptide search, these site-based results were then filtered further to allow comparison with the synthetic peptide known localisation key. Partial peptides and PSMs with the incorrect phosphorylation count compared to what was expected from the answer key, or other additional modifications, were removed. These remaining PSMs were then compared against the synthetic peptide answer key to determine if the phosphosites had been correctly identified. For analysis of the synthetic data only, results were ordered by the corresponding site localisation probability rather than the combined probability, since due to the small size of the search database, not all pipelines could produce accurate estimates of PSM probability.

The sites reported for each analysis method were ordered by combined probability and global FLR was estimated for every ranked site, from which we can then later apply a threshold at the lowest scoring site that delivers a desired global FLR (e.g. 1%, 5% or 10%), similar to the q-value approach for standard database searching. The global FLRs for all of the data sets were estimated using two methods – called Model FLR and Decoy FLR method. For the synthetic data set a third method was also used, called Answer Key method (comparing against the known phosphosite answer key)

### Model FLR method

Firstly, we estimated the global FLR using the combined probabilities (Global Model FLR). For TPP, we use the 1 – final probability, to give the local FLR (PEP) for each given site scored in a ranked list. The “Global Model FLR” is calculated as a running sum of the local FLR divided by the count of rows (Equation 1) i.e. the estimated frequency of false localisations at each row in a ranked list, divided by the total number of reported observations.

Equation 1 Global Model FLR:

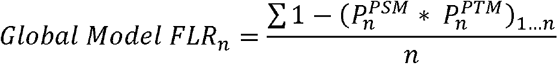

*Where n is the count of observations, P*^*PSM*^ *is the local FLR (PEP) for a given PSM identification and P*^*PTM*^ *is the local FLR for a given site localisation*.

### Decoy FLR method

We used the identification of phosphorylated decoy amino acids (e.g. worked example follows for Alanine, pA), as these are known to be false localisations and can therefore be used to estimate the FLR. The counts of the phosphorylated decoy amino acids were first normalised to allow comparison with true hits, modelling the *random* frequency one would expect incorrect sites to be assigned to target STY residues. The “STY:X ratio” was determined by dividing the total count of STY residues by the total count of the decoy amino acid residues, e.g. count Ser, Thr and Tyr / count Ala = *STY:A ratio*, within the set of PSMs with a scored phosphosite.

For every false localisation of the decoy amino acid, it would be expected to see *STY:A ratio X false localisations* within the target hits (STY). The count of phosphorylated amino acids is therefore multiplied by this ratio (to model the expected frequency of *random* wrong hits), then multiplied by 2 (to model the normalised frequency of *random* wrong assignments amongst both the decoy amino acid and the target amino acid) to arrive at a normalised false localisation count. This is relative conservative method to calculate global FLR, but without the correction to multiply by two, the use of a less frequent decoy amino acid would be insufficiently corrected for.

This normalised false localisation count is then divided by the total count of observations, at a given row in the ranked list, to obtain the Model (global) FLR estimate (Equation 2).

Equation 2 phosphorylated decoy amino acid FLR:

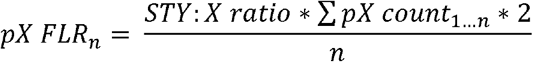

*Where pX count is the count of phosphorylated decoy amino acid and n is the count of observations at a given row in the ranked list*.

### Answer Key FLR

For the synthetic data set, we used the synthetic peptide false localisations in a similar way, the false localisation count (i.e. result not matching the answer key) was divided by the total count of sites to calculate the FLR (Equation 3).

Equation 3 Synthetic answer key FLR:

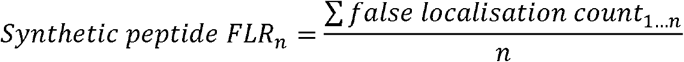

*Where n is the count of observations and false localisation count is the count of sites not matching the answer key in a given position in the ranked list*.

### Collapsing observations of a site across multiple PSMs

When summarising study results, it is desirable to “collapse” results where there are multiple PSMs supporting the same modification site down to a single row. There is no well-agreed method for collapsing data, although common practice when using collapsing multiple PSMs into individual reports for peptides is to use the maximum peptide score for ordering results, and disregarding the count of PSMs. The rationale for this simplification is that multiple PSMs reporting for the same peptide are not independent statistical tests and thus the same wrong answer can appear in multiple PSMs. As such, a simple method for ranking final “collapsed” results for sites is simply to take the maximum final probability. However, our own profiling of data sets suggests that this method is sub-optimal. Many of the high scoring decoy hits are supported by only a single PSMs, and so a collapse method that weights sites supported by a higher number of PSMs is more likely to be true than one supported by a single PSM (Supplementary Figure 2). This is an area of active development in our group to create a valid statistical model for multiple observations of a PTM site, which accurately models probabilities in this space. For this study, we use a relatively *ad hoc* method for collapsing multiple observations that attempts to balance maximum final probability and spectral counts. We took the maximum probability for a given site, derived from multiple PSMs, and binned into final probability values at 2 decimal places. We ranked via binned final probability, and then ranked within bins via the count of PSMs.

### Profiling distance distributions from real identifications to decoy amino acids

In order to compare between the decoy amino acids investigated, the distribution of amino acids around phosphorylation sites were compared. The phosphorylation sites obtained searching each database for phospho (STY) using TPP was first filtered for 5% model FLR. The minimum distance between an assumed correctly localised phosphorylated STY and the nearest candidate amino acid were compared, alongside the minimum distance for the nearest STY. Histograms were generated with the normalised frequencies of these distributions in order to compare between the selected decoy amino acids and STY.

### Profiling site probabilities for proximal amino acids

When analysing the results for different decoy amino acids we observed particular differences in the global FLR estimates for certain decoys (particularly pAla versus pGly) that could not be explained by distributions of amino acids in relation to confident target sites (above). We further explored these effects by calculating the average final probabilities for assumed correct sites with different amino acids in the −1 and +1 position relative to the site. The assumed correct sites were estimated as sites with combined probability ≥0.68. This threshold was calculated from the average minimum combined probability using a 5% FLR cut off for each of the decoy FLR estimations across all searches. These average probability distributions were calculated for the *Arabidopsis* and human data sets, from results of the TPP search with no decoy amino acid (pSTY), pAla decoy (pASTY), pLeu decoy (pLSTY) and pGly decoy (pGSTY).

## Results

### Analysis of synthetic data set PXD007058

The analysis setup aimed to determine whether global FLR within genuinely modifiable residues (target amino acids) could be estimated reliably by including in the search a “decoy” amino acid that is not modified. We tested six different amino acids to act as a decoy in parallel searches: glycine, leucine, alanine, glutamate, aspartate and proline, to determine what effect the selection of a particular decoy had on results obtained. The set of potential decoy amino acids were selected based on the following rationale: i) glycine, leucine, alanine – no evidence that they can be phosphorylated in any known biological system, all relatively frequent amino acids in most biological systems; ii) glutamate – infrequently phosphorylated [26] and not typically detectable as phosphorylated in most standard enrichment MS experiments, thus could be a plausible choice as a decoy; iii) aspartate and proline were chosen as expected to be deliberately poor choices of decoy amino acids, since there are known SP and SD phosphorylation motifs, which could bias estimates of global FLR. We expect a statistically reliable choice of amino acid should have a similar distance distribution from a phosphorylation site (STY) to another truly possible phosphorylation site (STY), under the theory that incorrect localisations are more likely to amino acids nearby in the sequence.

We first searched the PX007058 synthetic dataset, which allowed us to test the three different methods of FLR estimation against a known answer. The data set was searched using TPP and the global FLR of these was calculated using the decoy phosphorylated amino acid method for six different choices of decoy amino acid i.e. in six parallel searches (Figure 1a). As described in the Methods, the global “Decoy” FLR is estimated based on the counts of hits to the decoy amino acids in the ranked list of results, adjusted for the ratio of the counts of the decoy amino acid to the target amino acid in the modified peptides that have been considered. We also show the global FLR calculation for all three methods in Figures 1b–g, split by decoy amino acid choice i.e. i) *Answer key* - identifying false localisation by comparing to the known phosphosites (in this synthetic data set where the truly modified site is known), ii) Decoy amino acid method and iii) *Model* FLR i.e. based on summing local FLR calculated by the analysis software intrinsically (see Methods). The first observation we make is that the choice of decoy amino acid can have a substantial effect on the sensitivity (counts of assumed true sites), for a given estimated global FLR threshold (Table 2 and Figure 1). At 5% FLR, the lowest sensitivity is achieved with a Gly decoy (749 sites), versus the highest sensitivity with a Glu decoy (952 sites).

**Fig 1:**
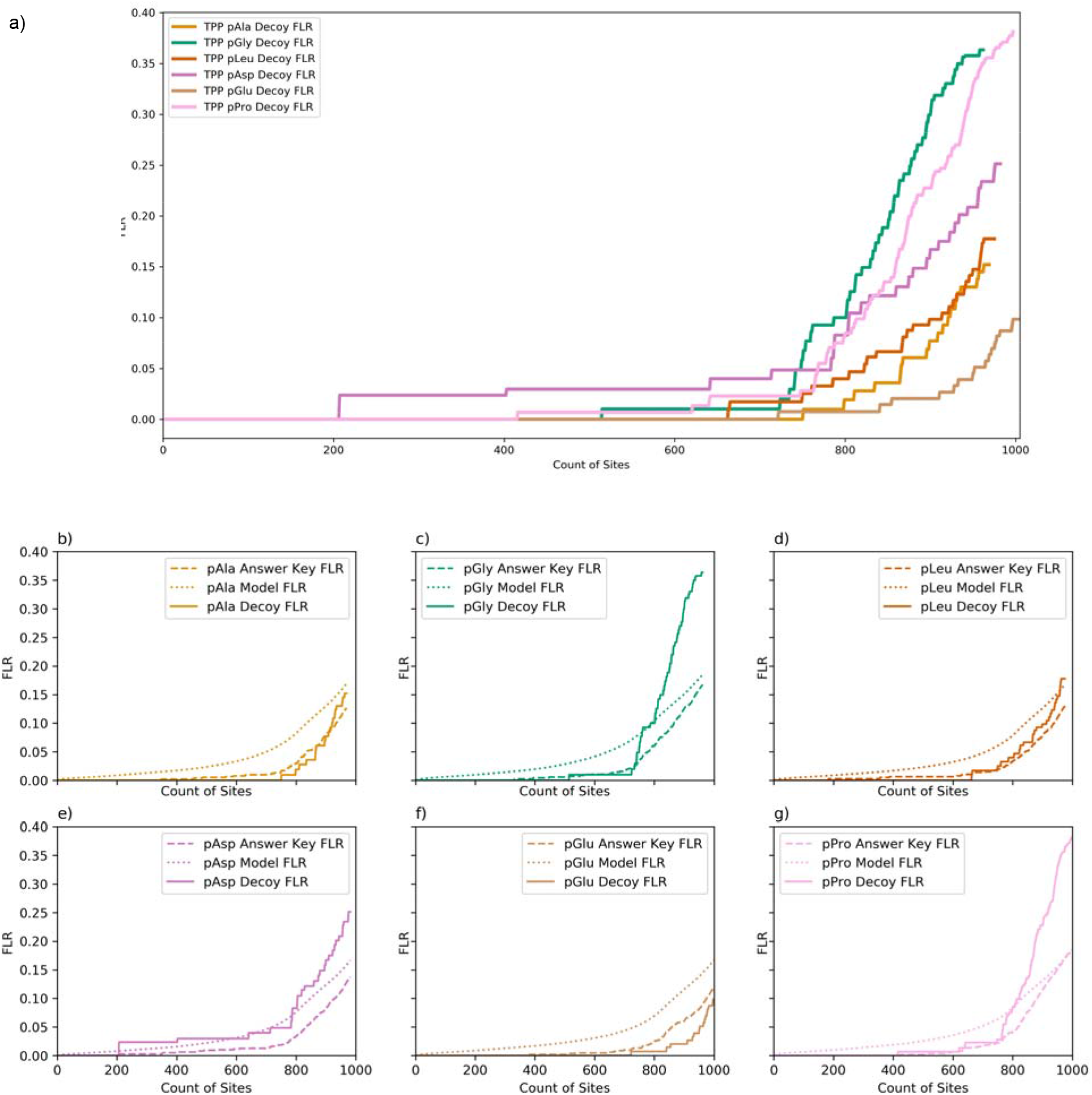
a) Comparison of FLR estimation searching PXD007058 (Synthetic data set) using different decoy amino acids: pAla, pGly, pLeu, pAsy, pGlu and pPro (TPP, fully tryptic, 1 %FDR) b-g) Comparison of FLR estimation methods searching PXD007058 for each of the different decoy amino acids (TPP, fully tryptic, 1% FDR). X-axis = count of sites, y-axis is global FLR estimated as q-values.

**Table 2:** Counts at pX FLR (calculated by the decoy method) for 1%, 5% and 10% using each FLR method, searching PXD007058 (Synthetic data set) (*TPP, fully tryptic, 1% FDR*)

|  | Count at 1% FLR |  |  | Count at 5% FLR |  |  | Count at 10% FLR |  |  |
| --- | --- | --- | --- | --- | --- | --- | --- | --- | --- |
|  | Answer Key | Model | Decoy | Answer Key | Model | Decoy | Answer Key | Model | Decoy |
| pAla | 702 | 226 | 799 | 835 | 700 | 866 | 935 | 836 | 921 |
| **pGly** | 571 | 203 | 515 | 778 | 645 | 749 | 866 | 791 | 787 |
| **pLeu** | 632 | 229 | 665 | 843 | 710 | 823 | 938 | 844 | 914 |
| **pAsp** | 535 | 258 | 207 | 834 | 721 | 784 | 929 | 849 | 805 |
| **pGlu** | 722 | 232 | 841 | 858 | 734 | 952 | 975 | 866 | n/a |
| **pPro** | 636 | 206 | 621 | 817 | 692 | 768 | 883 | 832 | 823 |

It can be seen that for pAla and pLeu decoys (Fig 1b and d), there is close agreement between the two “empirical” methods of estimating global FLR i.e. Answer Key and Decoy, with the Model giving more conservative estimates of FLR. The pGly and pPro methods (Fig 1c and g) have good agreement between the Answer Key and Decoy methods up to ~700 counts, and then the decoy method gives more conservative estimates (rising steeply), compared to the Answer Key method (and Model). The pAsp decoy method agrees well with the Model FLR but is more conservative than the Answer Key (Fig 1e). The pGlu decoy method is the least conservative, apparently underestimating global FLR compared to the answer key (Fig 1f). Overall, the matching of pAla and pLeu decoy FLR estimation to the answer key FLR gives some supporting evidence towards pAla and pLeu being appropriate choices for decoy amino acids. The Model FLR method is shown to be more conservative that the other “empirical” FLR methods in most cases, especially in the most important regions of the distribution i.e. up to 5% global FLR for example.

The estimates from Figure 1b–g and Table 2 demonstrate greater stability in the Model FLR and Answer Key FLR across different decoy amino acids i.e. there is less variation in sensitivity at a given estimated FLR. This is to be expected since comparing the six different searches, many of the errors in localisation are due to incorrect localisation to a target amino acid (which largely behave the same across the six searches).

On the same synthetic data set, we also compared the estimation methods across four different pipelines: TPP, PEAKS, MaxQuant and Mascot (Supplementary Fig 3, Supplementary Table 2). The PXD007058 synthetic data set was searched for pSTY and pAla and pLeu (for decoy comparison). The initial set of results from our analysis pipeline are the redundant identification of phosphorylation sites i.e. if multiple PSMs support the same site, these appear as multiple rows (not collapsed). In general, as noted in the Methods, our preference is to order these results by the *final probability* that a site has been observed (PSM probability X site localisation probability). The synthetic data set has a small database size and an overall small count of identifications, which makes it difficult to model PSM probability accurately. As such, for the synthetic data set only, we ordered results by the site localisation probability, having first accepted only PSMs with FDR < 1%.

FLR was calculated using the synthetic answer key and the decoy amino acid hits (Supplementary Fig 3). For all four pipelines tested, both the pAla and pLeu Decoy methods agree well with the results from the Answer Key FLR method across all three pipelines, demonstrating that our method with these amino acids gives reliable FLR estimates in a software-independent manner. There are differences in the total number of sites identified at a fixed FLR threshold, dependent on the pipeline applied. For this data set, TPP gives highest sensitivity, followed by PEAKS, Mascot/ptmRS and MaxQuant. However, our primary goal in this manuscript is not extensively to benchmark different pipelines, as there are choices of algorithm parameterisation, which need to be optimised and could affect conclusions, and thus we do not make any general conclusions about software performance for PTM analysis here.

### Biological data set analysis

#### PXD008355

To investigate the effect of using different decoy amino acids on different data sets, we compared the FLR estimations across the six different amino acids using two experimental data sets from *Arabidopsis thaliana* and Human. Figure 2 shows the decoy FLR comparisons searching the PXD008355 *Arabidopsis* data set with TPP. Here we can see a similar trend as previously seen in the synthetic data set FLR estimations with pGly and pPro giving most conservative performance at higher FLR values i.e. a steep rise in global FLR (Fig. 2a) and giving the lowest sensitivity at 10% global FLR, although there is a more complex picture at 1% and 5% FLR values (Table 4 and Fig. 2b). We assume that many studies will aim to threshold at 5% global FLR, here we observe lowest counts (of sites at 5% FLR) for the pAsp decoy, similar, intermediate counts for pGly, pGlu and pPro methods, and highest counts of sites (sensitivity) for pAla and pLeu decoys. One of the challenges with accurate FLR estimation is that there can be some high-scoring incorrect localisations, and their position in the ranked list can have significant implications on the count of sites at 1% FLR (Fig 2b and Table 3). We thus would not recommend generally thresholding at 1% global FLR, but instead applying a 5% FLR where the global FLR estimates are likely to be robust. There is additional discussion of these high-scoring false hits in the Supplementary materials (i), Supplementary Figure 4 and Supplementary Table 3. We also calculated the Model FLR for each decoy option, demonstrating good agreement at 5% FLR between the two methods (Decoy FLR versus Model FLR) for pAla, pLeu and pGly decoy options, but less good agreement for other decoys (Supplementary Figure 5).

**Fig 2:**
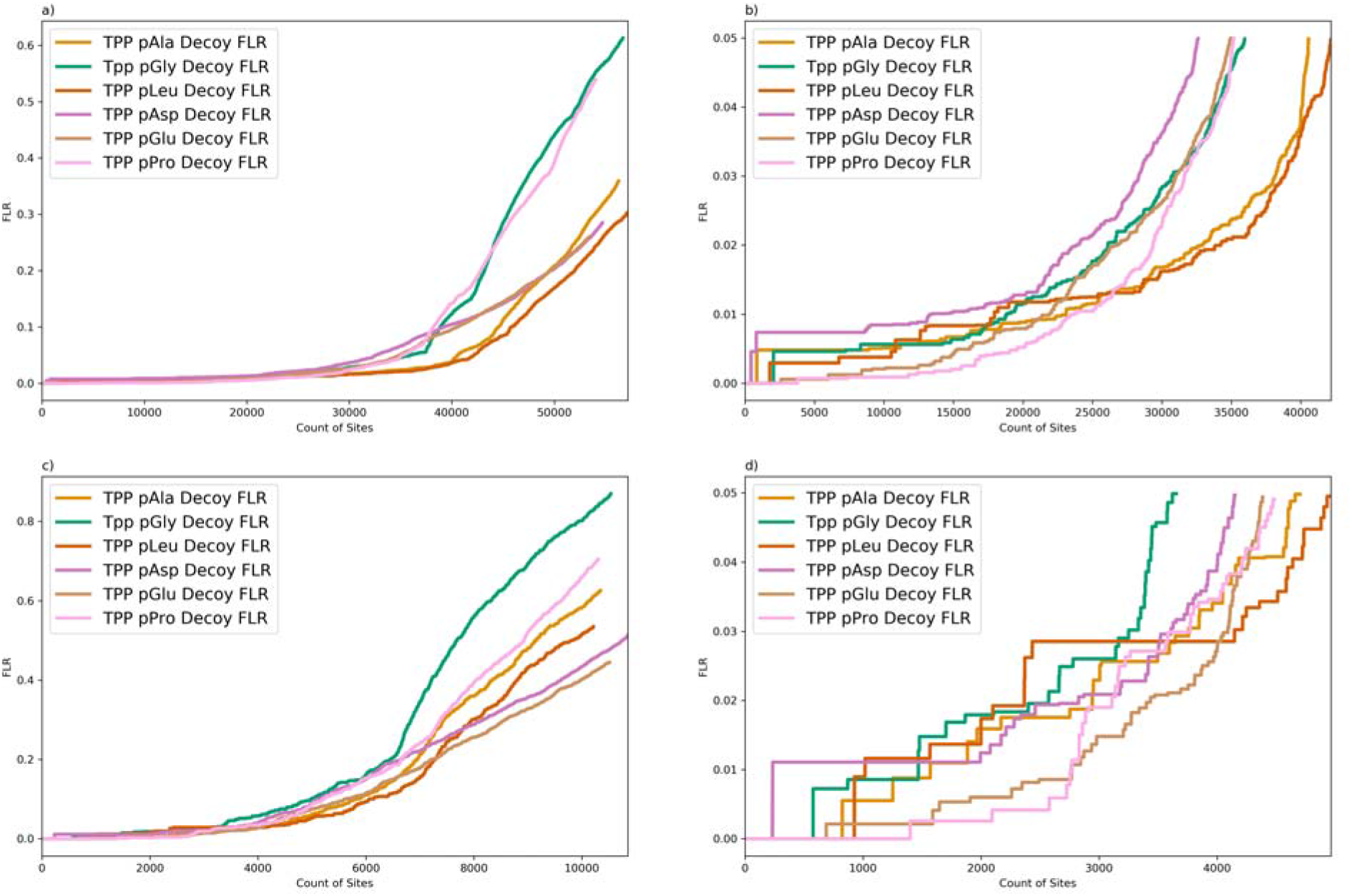
Comparison of FLR estimation searching PXD008355 (Arabidopsis data set) using different decoy amino acids: pAla, pGly, pLeu, pAsy, pGlu and pPro (TPP, fully tryptic, 1 %FDR) a) all PSMs, b) zoom 5%FLR all PSMs, c) collapsed by modified peptide, sorting by combined probability and count of supporting PSMs, d) collapsed by modified peptide, sorting by combined probability and count of supporting PSMs, zoom at 5%FLR. X-axis = count of sites, y-axis is global FLR estimated as q-values.

**Table 3:** Counts at pX FLR for 1%, 5% and 10% using each decoy method, searching PXD008355 (*Arabidopsis data set*) (*TPP, fully tryptic, 1% FDR*) *showing all PSMs (no collapse) and collapsing multiple PSMs to one row per modified peptide, sorting by combined probability and count of supporting PSMs*

|  | Count at 1% FLR |  | Count at 5% FLR |  | Count at 10% FLR |  |
| --- | --- | --- | --- | --- | --- | --- |
|  | No collapse | Collapse | No Collapse | Collapse | No Collapse | Collapse |
| **pAla** | 23104 | 1570 | 40541 | 4704 | 44556 | 5815 |
| **pGly** | 18872 | 1469 | 35939 | 3654 | 38885 | 4990 |
| **pLeu** | 17943 | 1017 | 42157 | 4964 | 45875 | 6068 |
| **pAsp** | 13490 | 234 | 32595 | 4151 | 39262 | 5130 |
| **pGlu** | 21923 | 2766 | 34949 | 4385 | 40532 | 5607 |
| **pPro** | 23696 | 2771 | 35170 | 4483 | 38226 | 5221 |

**Table 4:**
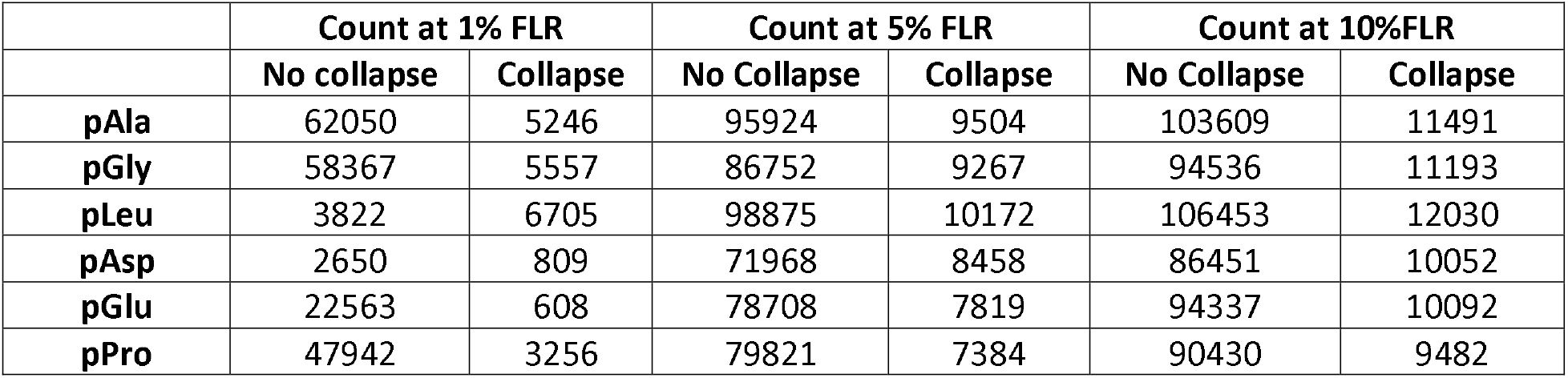
Counts at pX FLR for 1%, 5% and 10% using each decoy method, searching PXD000612 (human data set) (*TPP, fully tryptic, 1% FDR*) *showing all PSMs (no collapse) and collapsed to unique sites*.

|  | Count at 1% FLR |  | Count at 5% FLR |  | Count at 10%FLR |  |
| --- | --- | --- | --- | --- | --- | --- |
|  | No collapse | Collapse | No Collapse | Collapse | No Collapse | Collapse |
| **pAla** | 62050 | 5246 | 95924 | 9504 | 103609 | 11491 |
| **pGly** | 58367 | 5557 | 86752 | 9267 | 94536 | 11193 |
| **pLeu** | 3822 | 6705 | 98875 | 10172 | 106453 | 12030 |
| **pAsp** | 2650 | 809 | 71968 | 8458 | 86451 | 10052 |
| **pGlu** | 22563 | 608 | 78708 | 7819 | 94337 | 10092 |
| **pPro** | 47942 | 3256 | 79821 | 7384 | 90430 | 9482 |

We next explored data after collapsing multiple scores from different PSMs supporting the same site, taking the maximum probability for a given site for ranking results, along with the greatest number of supporting PSMs (see Methods). The results following this collapse step are shown in Figure 2c and d, demonstrating relatively similar trends, with considerable differences in sensitivity at 5% and 10% global FLR, with pAla and pLeu giving highest sensitivity at a 5% and 10% FLR. The statistical assumptions for the Model FLR do not hold after collapse, so this method was not used.

To further investigate the selection of decoy amino acid candidates, the minimum distance between an assumed correctly localised phosphorylated STY (<5% global FLR filtered) and the nearest candidate amino acid were compared, alongside the minimum distance for the nearest STY (Figure 3). The rationale for this comparison is that in a regular search not employing a decoy, if a phosphosite is wrongly localized, it will usually be to the nearest other STY residue than the correct site. We assume that a statistically reliable decoy amino acid will follow a similar (normalised) frequency distribution to the closest STY residue from correct hits. When comparing these distances in the *Arabidopsis* data set, it can be seen that Ala, Leu and Gly follow somewhat similar frequency distributions to proximal STY, particularly in the + positions (i.e. towards the C-terminus of the protein). Asp, Glu and Pro are all enriched at the +1 position relative to STY, which likely partially explains the higher FLR estimates observed for the same site counts in Table 3 and Figure 2, i.e. the pipeline wrongly assigns sites to ASp, Glu and Pro more frequently than it would be other target sites (STY). We also observed in Table 3 that using a glycine decoy gave relatively low sensitivity at 5% FLR, and steeply increasing FLR at higher site counts. The results for Gly in Figure 3 are thus are outlier with respective to Figure 2, as well as for the synthetic data set in Figure 1, in which we observed lowest site count at 5% FLR for estimates using a Gly decoy. Our starting expectation was that Ala, Leu and Gly would all make reliable choices as decoy amino acids, and thus we also conducted an analysis of amino acid frequencies to attempt to explain the differences seen for Gly, results are shown in Supplementary material (ii) and Supplementary Table 4. Gly and Ala have similar frequencies of observations in phosphopeptides, so this also does not explain the disparity. We further explore this phenomenon below.

**Fig 3:**
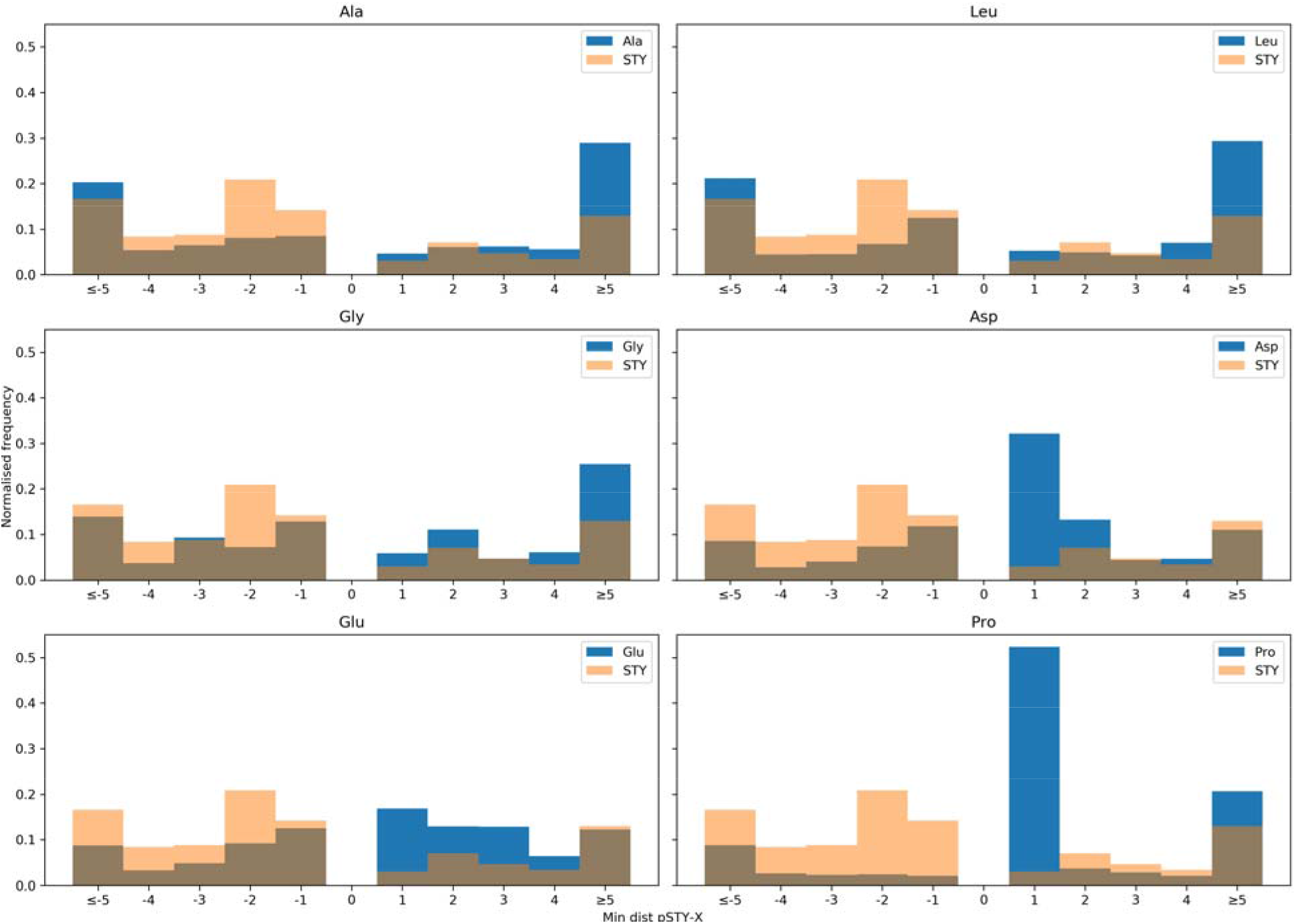
Comparison of minimum distance between phosphorylated STY and the nearest target amino acid (Ala, Leu, Gly, Asp, Glu and Pro), compared to the STY distribution, searching PXD008355 (Arabidopsis data set).

Similar to the synthetic data set, a comparison between FLR estimates was made across the different pipelines: TPP, PEAKS, MaxQuant and Mascot/ptmRS. The PXD008355 *Arabidopsis* data set was searched with an Ala decoy, as well as a Leu and Gly decoy, and FLR estimations were calculated in the same way as before (Supplementary Figure 6, Supplementary Table 5). In general, highest sensitivity is achieved by TPP and Mascot/ptmRS, whereas there are high-scoring decoy (amino acid) hits in the other two pipelines that lead to much lower sensitivity at a given FLR cut-off. For TPP and Mascot/ptmRS pipelines, the results from estimation with the three decoys are largely reproducible i.e. pAla and pLeu gives highest (and similar) counts of sites at a given FLR, whereas pGly gives a lower count of sites at the same FLR threshold.

In this approach, sites are ordered by final probability (PSM probability * PTM probability). An alternative approach commonly used in the field is to threshold first at say <1% FDR for PSMs or peptides, and then order purely by PTM localisation score or probability. We tested a similar approach to see what effect there is on sensitivity at a given FLR for the pAla results (Supplementary Fig 7). Whilst ordering site localisations by PTM probability only rather than the combined PSM*PTM probability, we can see that there is lower sensitivity at 1% FLR for the PTM probability option, and almost identical sensitivity between the two options at 5% and 10% FLR (Supplementary Table 6). We therefore conclude that it is slightly superior to model both the probability that a given PSM is correct, as well as that the PTM has been correctly localised to give the best ordering of results, particularly for those highest scoring around 1% global FLR.

#### PXD000612

Given that we see consistent trends for the synthetic data set and *Arabidopsis* data set in terms of comparing decoys across different pipelines, for the final validation, we focus only on the use of TPP on one further validation set from a different species (human). We would expect some different phosphorylation motifs comparing an animal species to a plant species, which could affect decoy amino acid performance. Figure 4 and Table 4 illustrate the FLR comparison using the different decoy amino acids, searching the PXD000612 human data set. A similar trend can be seen here as in the *Arabidopsis* data set with pAla and pLeu giving the highest site counts at 5% FLR and pAsp giving lowest site count. On the zoomed plot (<5% FLR, Figure 4b), the same issue as for data set PXD008355 can be observed, with unstable decoy estimation at low counts due to a random factor from a few high-scoring decoys (FLR < 1%). For this data set, there is also good agreement between the Model FLR and the Decoy FLR for most amino acids except pGlu, where the Model FLR tends to be more conservative than Decoy FLR (Supplementary Figure 8). We also show the data after collapsing multiple PSMs reporting on the same site (Fig 4c and d), giving similar trends in sensitivity at fixed FLR thresholds as for data without collapse.

**Fig 4:**
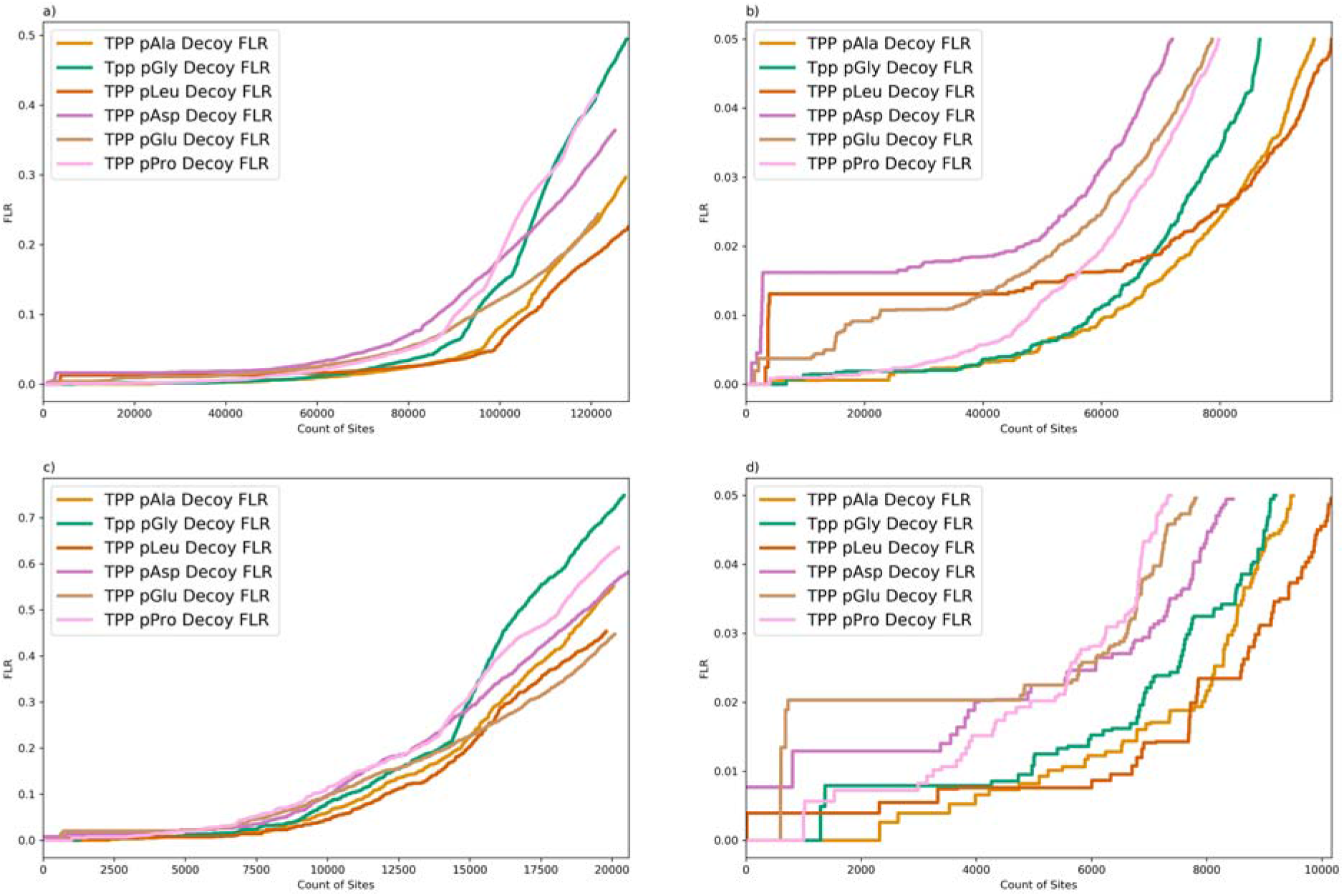
Comparison of Decoy FLR estimation searching PXD000612 (human data set) using different decoy amino acids: pAla, pGly, pLeu, pAsy, pGlu and pPro. (TPP, fully tryptic, 1 %FDR) a) no collapse, all sites shown, b) zoom on 5% FLR, c) collapse to unique sites, d) zoom on <5 FLR for data collapsed to unique sites. X-axis is count of sites, y-axis is global FLR, estimated as q-values by the Decoy method.

The distance between the phosphorylated STY and the nearest candidate amino acid were again compared (Supplementary Figure 9) to further investigate the effect of decoy amino acid choice and to examine how the distributions differ between the different data sets. When comparing these distances in the Human data set, a similar pattern is seen to that of the *Arabidopsis* data set. It can be seen that Ala and Leu again follow a somewhat similar frequency distribution to proximal STY residues, again particularly in the positive direction. Asp and Pro are again enriched at the +1 position relative to STY, which would be expected. Gly is also seen to follow a similar distribution to STY residues and therefore would be expected to be a reliable decoy amino acid, based on this measure. However, looking at the FLR comparisons seen in Fig 4, Gly can be seen to give more conservative FLR estimation (or lower site counts at 5% FLR for example than Ala or Leu), as was also seen in the *Arabidopsis* data set.

We next explored whether particular amino acids in proximity to true phosphorylation sites cause results to change. We plotted the average final probabilities for the *Arabidopsis* data set searched via the TPP pipeline, split according to the amino acid in the −1 (Fig. 5) and +1 (Supplementary Figure 10) position relative to the assumed correct phosphorylation site, for the data set searched with no decoy pSTY, pAla decoy, pLeu decoy and pGly decoy. In the search with no decoy, there is a particularly striking trend that sites have a lower probability when the −1 amino acid is Ser, Thr or Tyr. This occurs because the site localisation algorithm (PTMProphet) has fewer ions available to discriminate the correct from incorrect localisation. In the pAla and pLeu results, we see that Ala and Leu in the −1 position, cause sites to have a similar reduction in final probability as Ser, Thr and Tyr in the no decoy search i.e. final probability shifts from around ~0.96 to ~0.91 (Ala and Leu decoy). We interpret this to mean that they behave as statistically “good decoys” i.e. when they are present in the −1 position relative to a true site, they behave in a similar manner to STY residues. In the pGly data, there is a much larger drop off in final probabilities when G is in the −1 position (~0.96 to~0.87), meaning that (most commonly) Gly-pSer sites are scored less well that phosphoserines preceded by other amino acids, and excessive probability space is being distributed to the pGly-Ser hypotheses. In the results we observe that around 47% (*Arabidopsis* data) and 35% (human data) of the high scoring pGly decoys have pGly-Ser motif. We see similar trends in the human data set (supplementary Figures 11 and 12). It is unclear why this particular amino acid combination causes a problem for PTM localisation, but we hypothesise that the Gly-pSer bond is perhaps particularly stable during fragmentation and hence a discriminating y ion terminating with pSer is less commonly observed. We thus conclude that based on what we have observed that pGly is not an ideal choice for a decoy amino acid.

**Fig 5:**
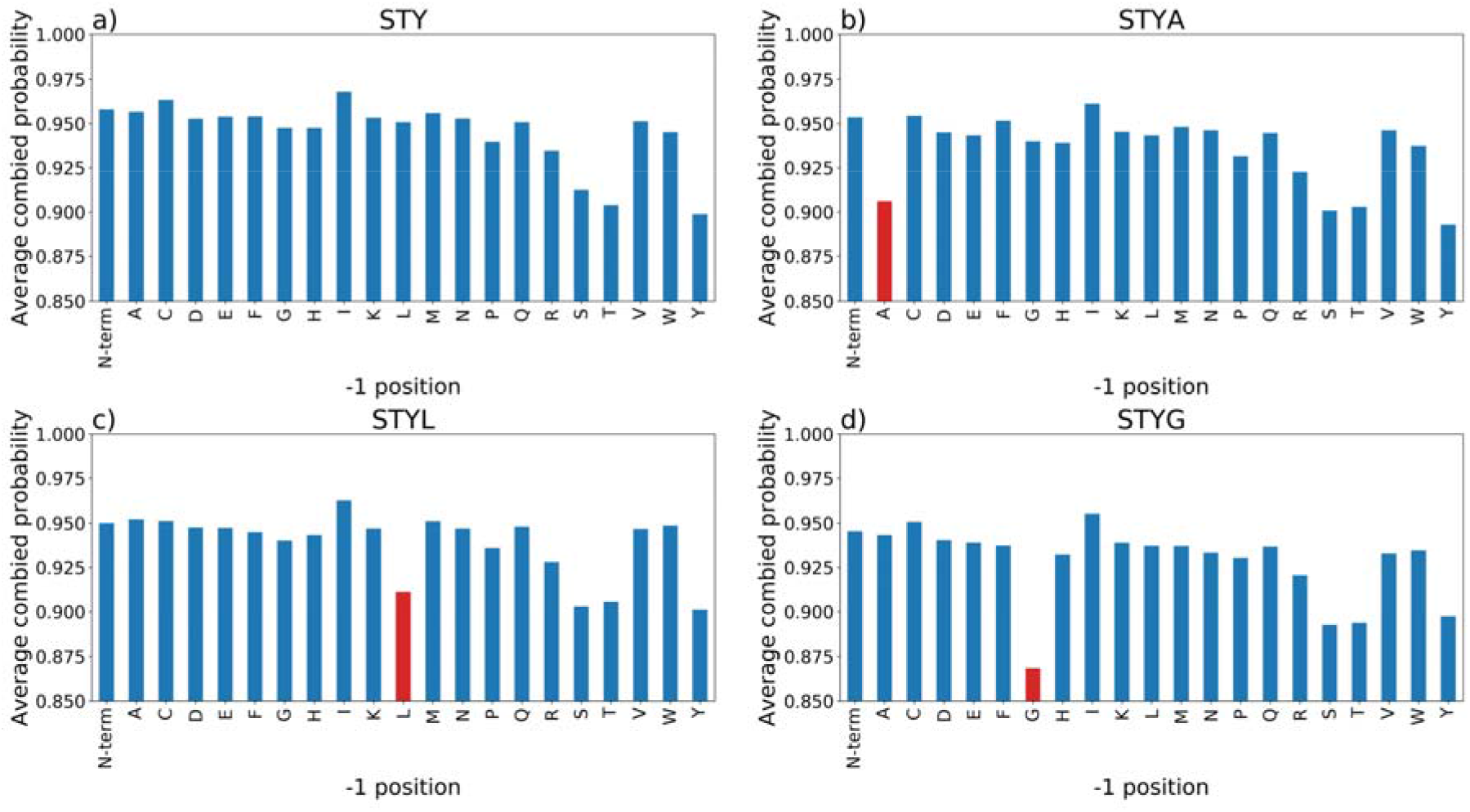
Comparison of averaged final site probabilities for all peptides (final probability ≥0.68 split by amino acid in the −1 positions for the PXD008355 (Arabidopsis data set) a) STY (no decoy), b) STY with Ala decoy, c) STY with Leu decoy and d) STY Gly decoy.

## Discussion

In the proteomics field, prior to the widespread adoption of decoy database searching, there was a general problem with false positive results in the literature, as labs vied to report the largest number of peptides and proteins, without control of FDR. It is now accepted that all proteomics studies should control for FDR at an appropriately conservative level, e.g. < 1% FDR at the protein-level for protein-centric studies. It has long been recognised that a similar problem exists with reporting PTM sites. The accurate discovery of a site can be crucially important for downstream interpretation, since the identity of the residue (STY for canonical phosphorylation) and the proximal amino acids govern understanding of the kinase and phosphatase that regulate it. Given the interest in understanding phosphorylation (and other PTM) sites in most human diseases, adequate control of false reporting is crucial. It has recently reported that >80% of reported sites in a popular phosphorylation database are estimated to be false positives [27]. This has likely resulted due to studies using overly weak FLR thresholds in publications, and results then get deposited in databases. Correct identifications tend to be reported from multiple studies, whereas *random* wrong site identifications tend to be seen only one or twice and thus over time, database-level FLR creeps up.

This study is, to our knowledge, the most detailed attempt to understand how best to estimate global FLR using decoy amino acids. We compare the method against the use of a statistical model, based on summing local FLR values, and results agree reasonably, but not perfectly well. We also demonstrate that the selection of a particular amino acid, even when correcting for the frequency of that amino acid in the results, does influence results more than would be desirable. We believe that our results back up that either pAla or pLeu make appropriate decoys based on their similar frequencies proximal to real phosphorylation sites (in a test case from humans and a model plant), as compared to target amino acids STY. The results for pAla and pLeu decoys also agree well with the Model FLR (for the large data sets) and the Answer Key FLR, for the synthetic data set. We have a slight preference to use pAla as a decoy going forward, since there is a slight risk of confusion between Leu and Ile amino acids, which often cannot be distinguished by MS. In rare cases where there are two peptides in the database, differing only by Ile/Leu, errors or inconsistencies in decoy FLR estimation could be introduced.

From the TPP pipeline, using iProphet and PTMProphet, it is possible in theory to use either the Model FLR or the Decoy FLR for thresholding final results. As noted above, performing a 1% global FLR threshold may be unstable (based on the Decoy FLR method), depending on the chance appearance of a few decoys high on the ranked list. If control at this level is required, the Model FLR would thus be preferred. For a less conservative threshold, say 5% FLR, then we believe that thresholding using the pAla Decoy FLR method should be recommended. The rationale is that this method can be straightforwardly applied using any combination of tools, and is simple to interpret. Most other pipelines in current use do not report accurate PEP values for PSMs and for site localisation, allowing Model FLR to be calculated reliably. We also recommend that the scores per site (final probability in the case of TPP-produced data) and the pAla “identifications” get carried forward and reported. This allows for the potential for meta-analyses and database submissions to estimate the resulting global FLR once multiple data sets have been combined.

We acknowledge that there is a slight downside to searching with a decoy amino acid, in that the search spaces for PSM identification and PTM localisation are both increased, leading to a potential loss in sensitivity. In our analyses, Ala residues are present at less than 1/3 the total frequency of STY residues, leading to a relatively modest increase in search spaces, of say 30%. We also suggest that the proteomics field has generally accepted that doubling the PSM search database (and search time) through the inclusion of decoys is an acceptable trade-off for gaining the ability to estimate global FDR straightforwardly and transparently. While we have presented results for pAla as a decoy for phosphorylation studies, we also suggest that modified Ala could also be an appropriate decoy for other modification types, such as Lys modifications acetylation, methylation, ubiquitination and SUMOylation etc, although we have not yet profiled the amino acid distributions sufficiently to conclude that Ala is more suitable than other amino acids in these cases.

## Conclusions

We have assessed six different amino acids for their ability to act as suitable decoy amino acids for the estimation of global FLR in phosphoproteomics studies. We have analysed three data sets, one synthetic with a known answer and two biological-sample data sets. We conclude that either Ala or Leu make appropriate decoys, and give reliable estimates of FLR above 1% FLR. Below 1% FLR, estimates can be unstable due to a few *random* high-scoring decoys. We demonstrate that the decoy-based FLR gives similar estimates to a modelled FLR for Ala and Leu decoys, based on summing local FLR values per site, and based on the answer key for the synthetic data set. We recommend that phosphoproteomics investigators should adopt the “pAla” decoy going forward i.e. the *pASTY method*, and report sites with appropriate global FLR control.

## Acknowledgements

We are grateful for funding from BBSRC/NSF that supported this work [BB/S017054/1, BB/S01781X/1]. JAV and MM would also like to acknowledge EMBL core funding. EWD acknowledges funding by NSF grant DBI-1933311 and NIH grants R01GM087221 and R24GM127667.

