## Supplementary materials for "A method for independent estimation of false localisation rate for phosphoproteomics"

*Supplementary Table 1: Search parameters for each data set, using PEAKS, MaxQuant and Mascot/ptmRS pipelines. *PEAKS only, **MaxQuant only*

|  | **PXD007058 (Synthetic data set)** | **PXD008355 (*Arabidopsis* data set)** |
| --- | --- | --- |
| **Peptide Mass Tolerance** | 20.0 ppm | 10 ppm |
| **Fragment Bin Tolerance** | 0.02 Da | 0.02 Da |
| **Digest Mode** | Tryptic | Tryptic |
| **Max Missed Cleavages** | 2 (4*) | 2 |
| **Fixed Mods** | Carbamidomethylation (C) | Carbamidomethylation (C) |
| **Variable Mods** | Oxidation (MWP) (Oxidation (M)*)  Phospho (STYX)  Pyrophospho (STY),  N-terminal acetylation**  Ammonia loss (QC)**  Pyro-Glu (EQ on the N-terminus)**  Deamination (NQ)** | Oxidation (M)  Phospho (STYX)  N-terminal acetylation  Ammonia loss (QC)  Pyro-Glu (EQ on the N-terminus)  Deamination (NQ) |
| **Max Variable PTMs** | 3 (5*) | 3 |

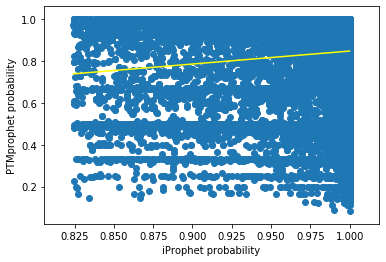

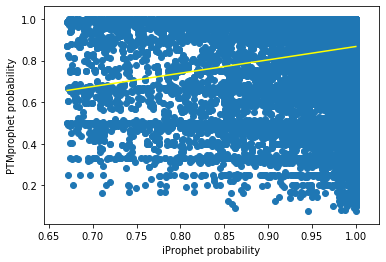

*Supplementary Fig 1: log10 PTM score vs Score. a)PXD008355 (Arabidopsis data set) R^2^=0.00740, b)PXD000612 (Human dataset) R^2^=0.0132.*

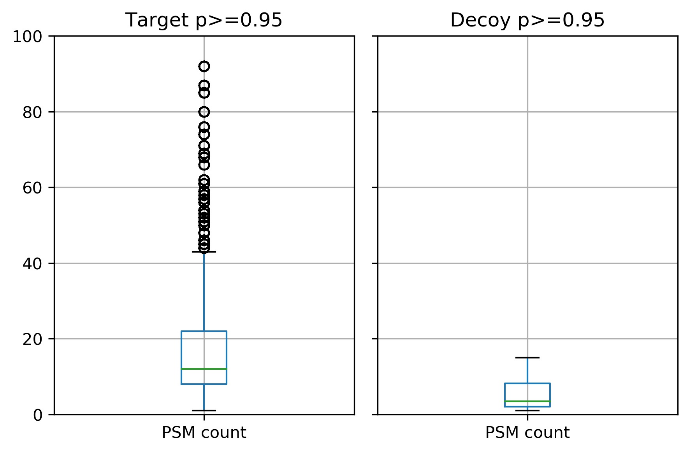

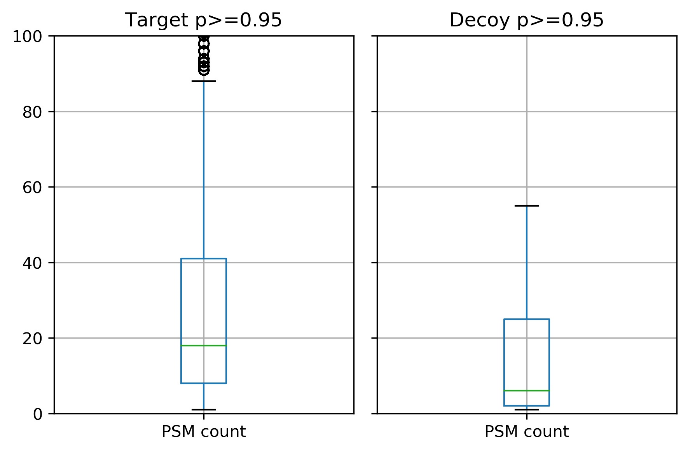

*Supplementary Fig 2: Boxplots of PSMs supporting targets and decoys with final probability >0.95 (pSTYA, 1%FDR). a)PXD008355 (Arabidopsis data set), b)PXD000612. (Human data set)*

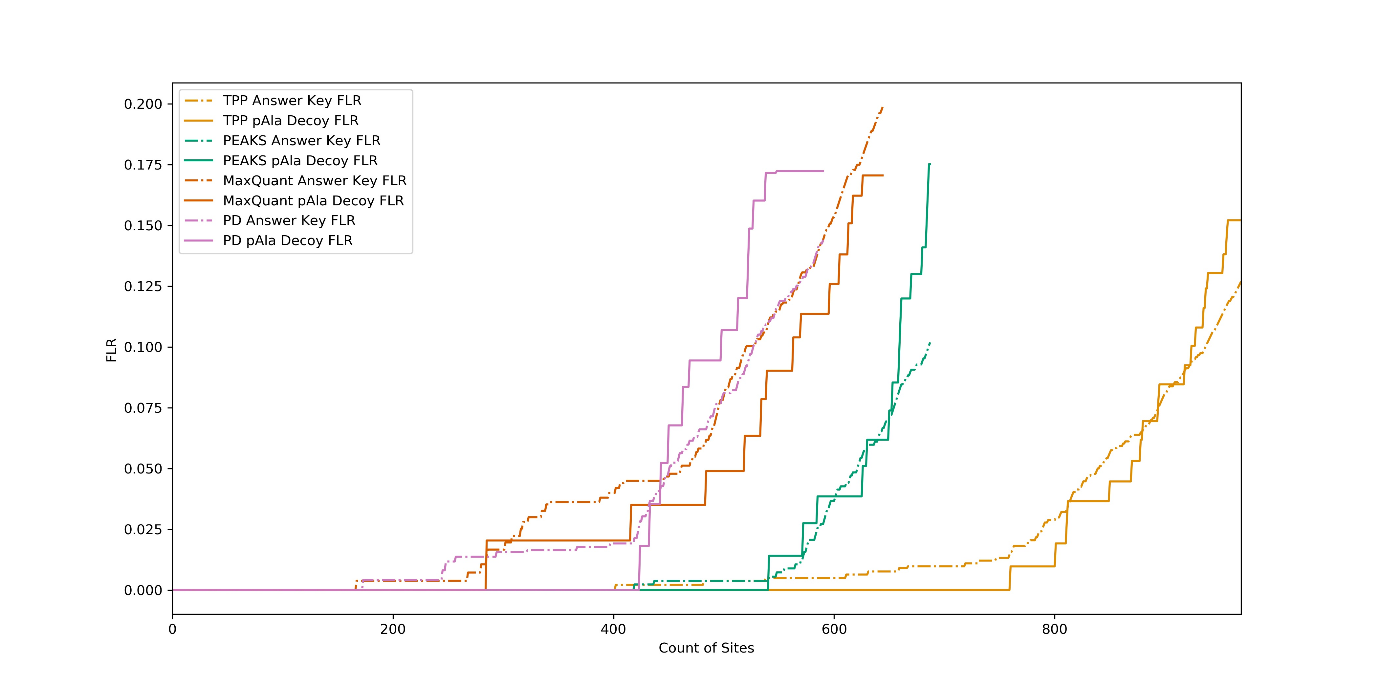

a)

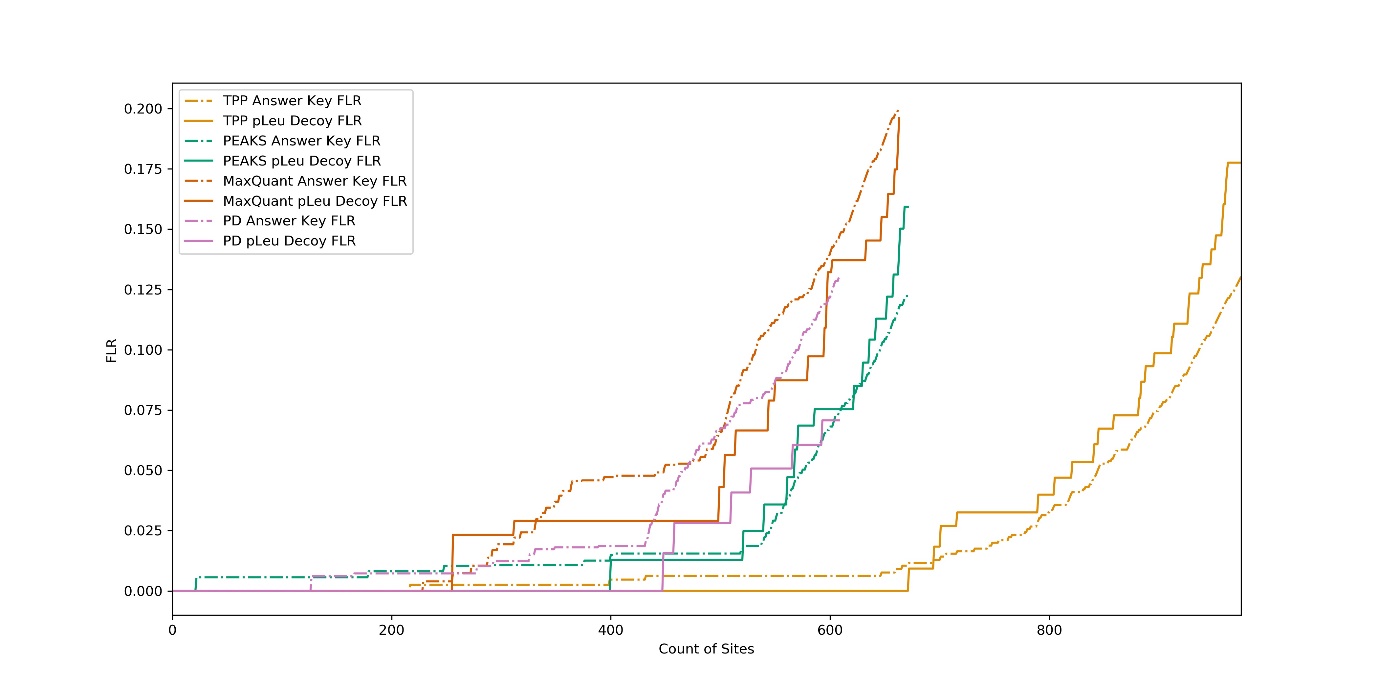

b)

*Supplementary Fig 3: Comparison of pX FLR estimation searching PXD007058 (Synthetic data set) using different pipelines: TPP, PEAKS, MaxQuant and Mascot/ptmRS (PD). PSMs are ordered by PTM probability (Fully tryptic, 1 % FDR). a) pAla decoy, b) pLeu decoy.*

*Supplementary Table 2: Comparison of pAla/pLeu Decoy FLR estimation searching PXD007058 (Synthetic data set) using different pipelines: TPP, PEAKS, MaxQuant and Mascot/ptmRS (PD). PSMs are ordered by PTM probability (fully tryptic, 1% FDR).*

|  | **Count at 1% FLR** | | **Count at 5% FLR** | | **Count at 10% FLR** | |
| --- | --- | --- | --- | --- | --- | --- |
|  | **pAla** | **pLeu** | **pAla** | **pLeu** | **pAla** | **pLeu** |
| **TPP** | 801 | 695 | 870 | 821 | 924 | 912 |
| **PEAKS** | 541 | 400 | 626 | 568 | 660 | 636 |
| **MaxQuant** | 285 | 256 | 519 | 504 | 563 | 595 |
| **Mascot/ptmRS** | 424 | 448 | 443 | 528 | 498 | 609 |

i) **Investigating high-scoring false hits**

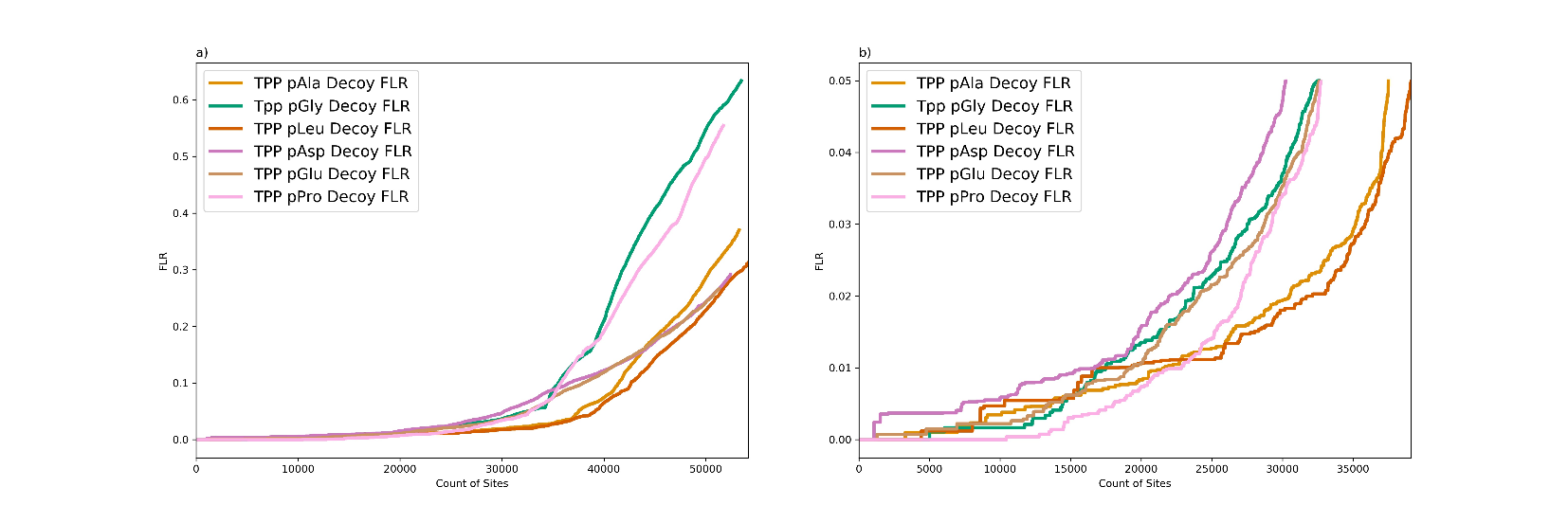
When searching the PXD008355 *Arabidopsis* database with TPP using different decoy amino acids, multiple high scoring false localisations could be seen. When these were investigated further, it was found that these wrong hits contained the same number of potential phosphosites as identified phosphosites. These wrong hits can therefore be categorised as “no-choice” PSMs as there is no choice for localisation, which may impact on the scoring of these positions. This may indicate that the search engine and PSM scoring is producing over-confident estimates of probability. These “no-choice” hits were removed and the FLR estimations recalculated, resulting in an improvement in the FLR estimations for each method.

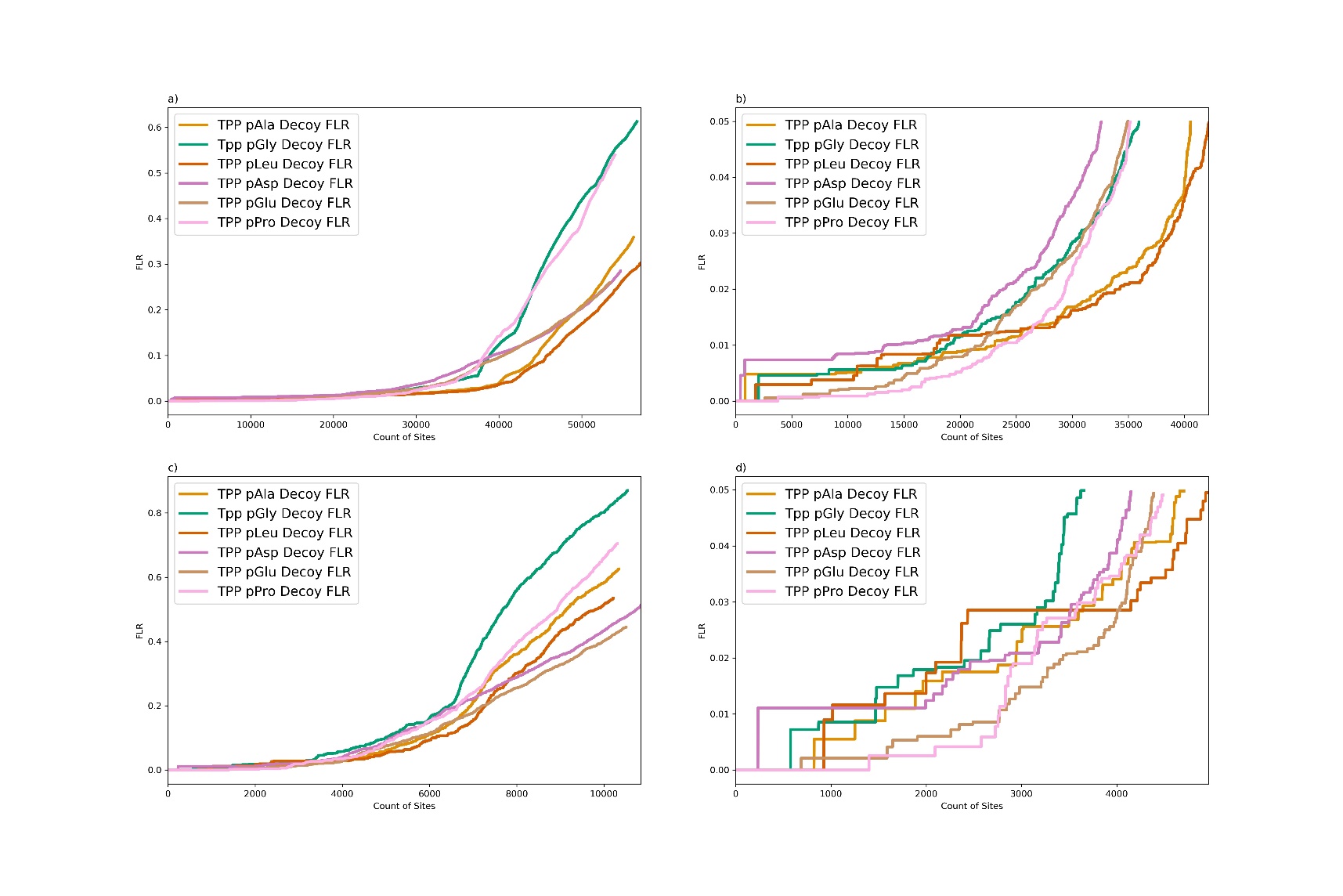
*Supplementary Figure 4: Comparison of FLR estimation searching PXD008355 (Arabidopsis data set) using different decoy amino acids: pAla, pGly, pLeu, pAsp, pGlu and pPro (TPP, fully tryptic, 1 %FDR). a) all PSMs, b) FLR ≤ 0.05 , c) all PSMs with “no-choice” hits removed, d) FLR ≤ 0.05 with “no-choice” hits removed. Figs a & b are shown in the main body of the manuscript and are repeated here for comparison.*

*Supplementary Table 3: Counts of sites at pX Decoy FLR for 1%, 5% and 10% threshold using each decoy amino acid: pAla, pGly, pLeu, pAsp, pGlu and pPro, searching PXD008355 (Arabidopsis data set) with “no-choice” hits removed (fully tryptic, 1% FDR).*

|  | **Count at 1% FLR** | **Count at 5% FLR** | **Count at 10% FLR** |
| --- | --- | --- | --- |
| **pAla** | 21689 | 37502 | 41411 |
| **pGly** | 17459 | 32570 | 35515 |
| **pLeu** | 16681 | 39118 | 42650 |
| **pAsp** | 16714 | 30236 | 36566 |
| **pGlu** | 19331 | 32584 | 37963 |
| **pPro** | 22839 | 32716 | 35743 |

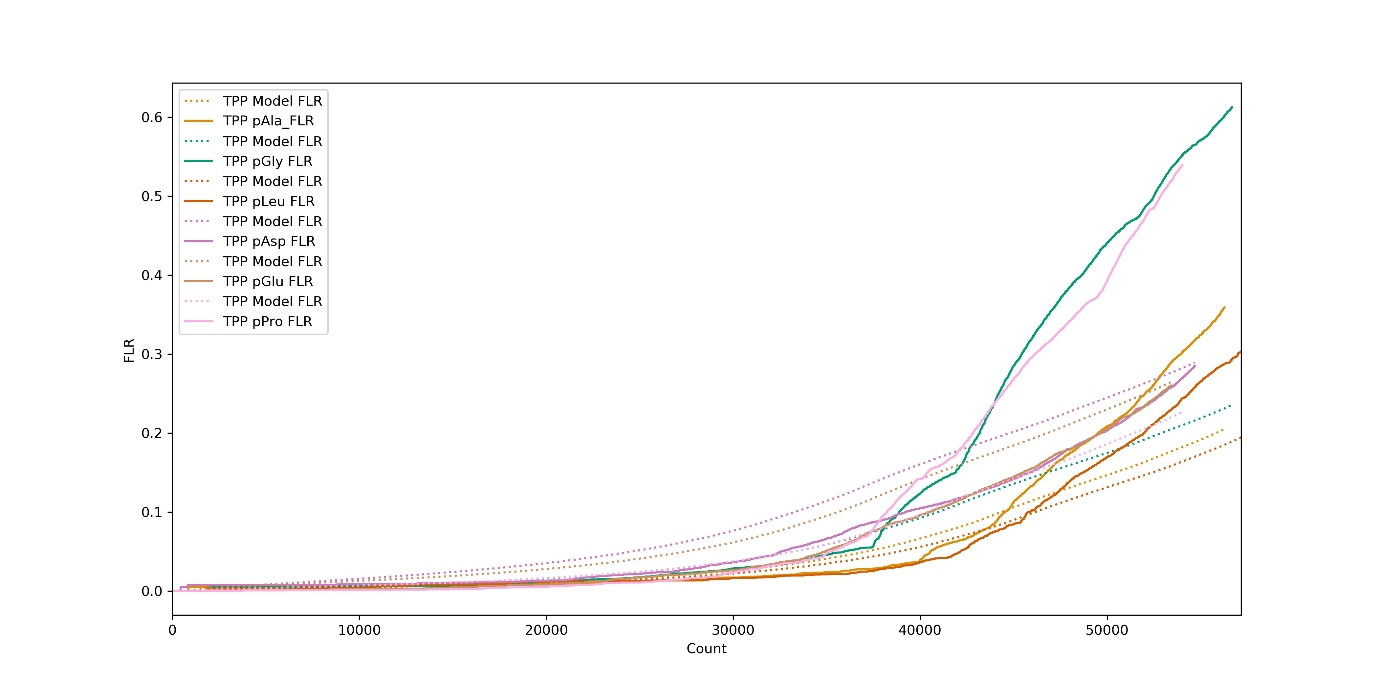

*Supplementary Fig 5: Comparison of pX Decoy FLR and Model FLR estimation searching PXD008355 (Arabidopsis data set) for pAla, pLeu, pGly, pAsp, pGlu and pPro (Fully tryptic, 1 %FDR).*

ii) **Amino acid frequency analysis**

In order to try to determine the cause of the differences seen with glycine, the amino acid frequencies of the decoys used were compared across the identified peptides, phosphopeptides and search databases. It could be seen that the frequencies across peptides and phosphopeptides of Ala and Gly, compared to STY were similar across the data sets. Although differences were seen between the search databases and peptide/phosphopeptide frequencies, the database frequencies showed fairly similar frequencies across all decoy amino acids. The analysis of the amino acid frequencies was therefore unable to determine an explanation for Gly being seen as an outlier.

*Supplementary Table 4: Comparison of amino acid frequency ratios between STY and the decoy amino acid for the identified peptides, identified phosphopeptides and the search database*

|  | **PXD007058**  **(Synthetic data set)** | | | **PXD008355**  ***(Arabidopsis data set)*** | | | **PXD000612**  **(Human data set)** | | |
| --- | --- | --- | --- | --- | --- | --- | --- | --- | --- |
|  | **Peptides**  **STY:X** | **Phosphopeptides**  **STY:X** | **Database**  **STY:X** | **Peptides**  **STY:X** | **Phosphopeptides**  **STY:X** | **Database**  **STY:X** | **Peptides**  **STY:X** | **Phosphopeptides**  **STY:X** | **Database**  **STY:X** |
| **Ala** | 3.67 | 3.57 | 4.22 | 2.57 | 3.49 | 2.72 | 3.04 | 3.32 | 2.33 |
| **Gly** | 3.74 | 3.88 | 3.66 | 2.40 | 3.12 | 1.79 | 2.98 | 3.36 | 1.64 |
| **Leu** | 3.40 | 3.37 | 3.45 | 3.29 | 4.55 | 2.68 | 4.30 | 4.39 | 2.49 |
| **Asp** | 4.23 | 4.20 | 5.74 | 1.60 | 1.88 | 3.14 | 2.90 | 2.87 | 3.45 |
| **Glu** | 2.42 | 2.44 | 3.62 | 1.35 | 1.74 | 2.50 | 2.05 | 2.07 | 2.30 |
| **Pro** | 1.82 | 1.84 | 2.45 | 2.89 | 2.60 | 3.58 | 2.02 | 1.98 | 2.59 |

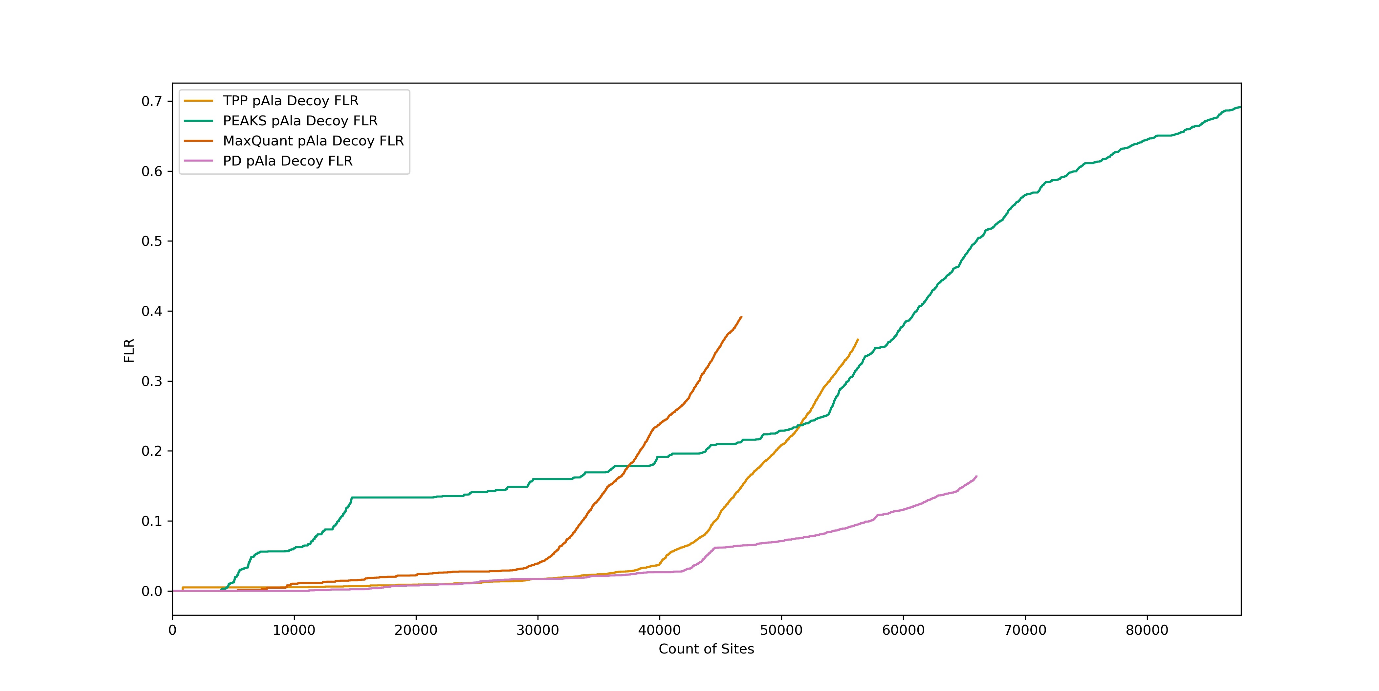

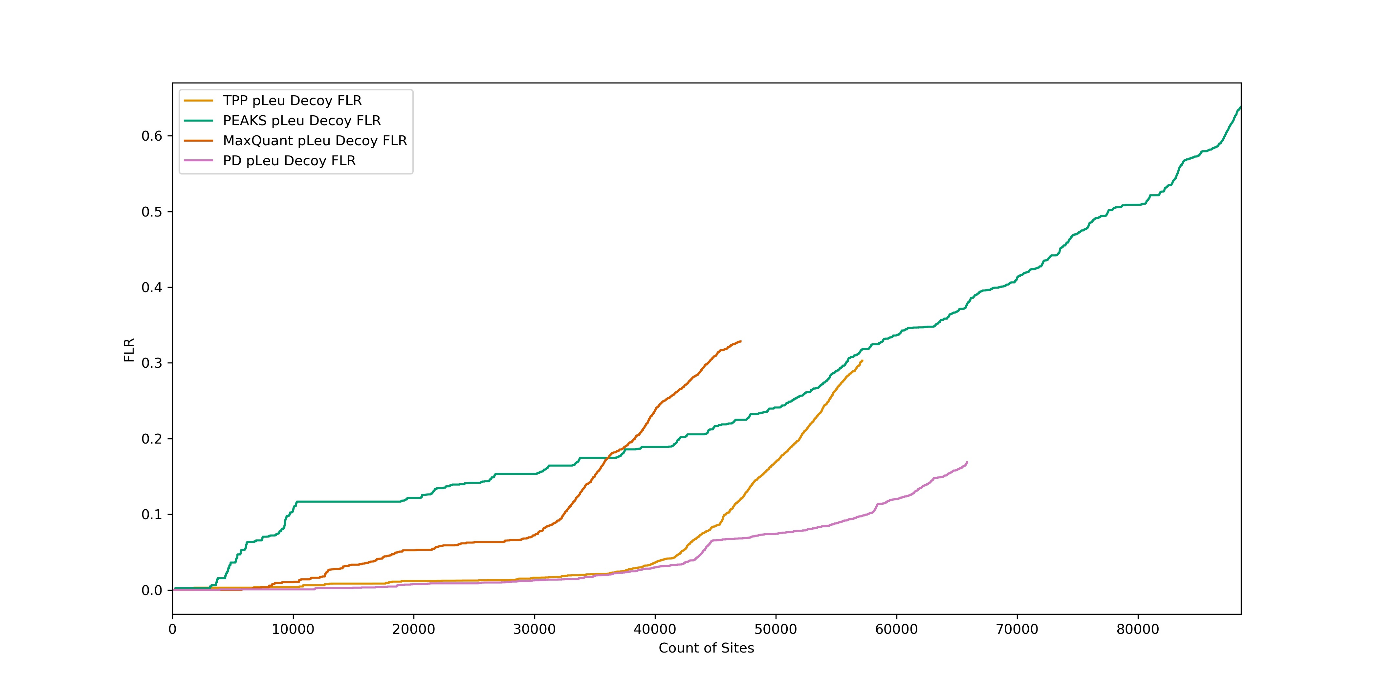

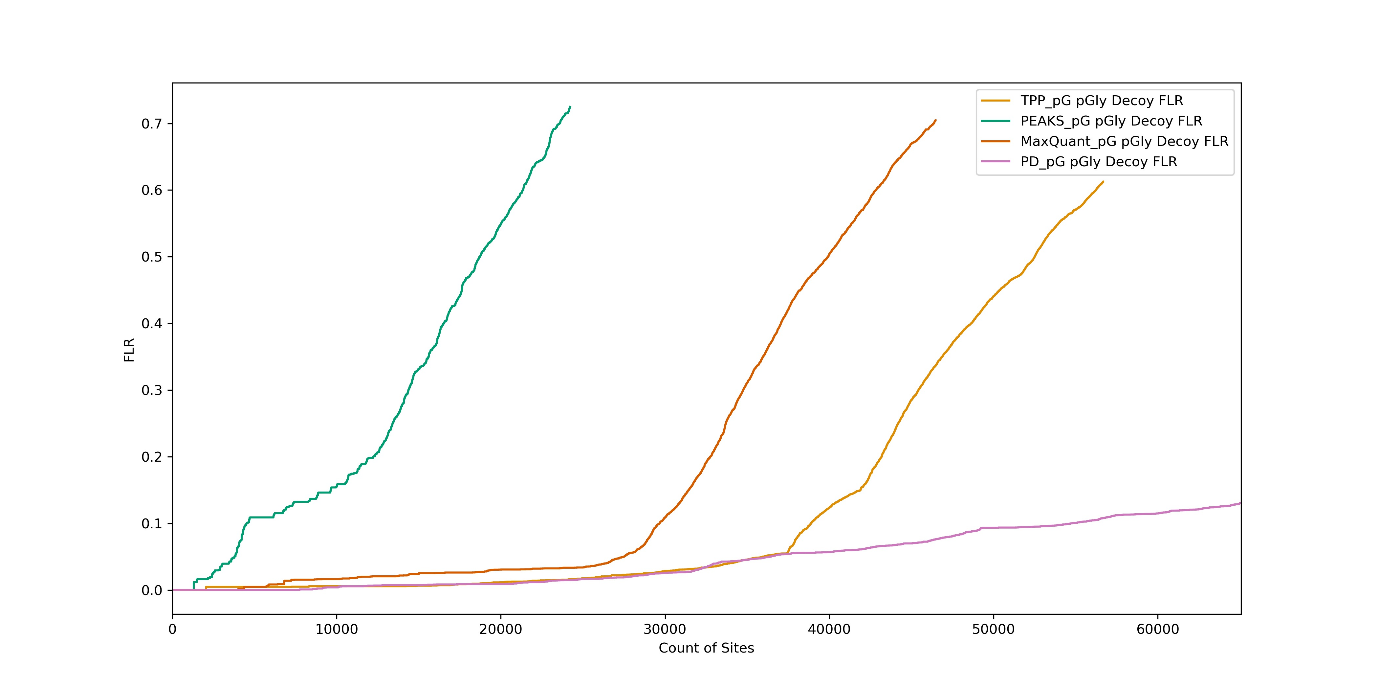

*Supplementary Fig 6: Comparison of a) pAla, b) pLeu and c) pGly Decoy FLR estimation searching PXD008355 (Arabidopsis data set) using different pipelines: TPP, PEAKS, MaxQuant, Mascot/ptmRS (PD) (Fully tryptic, 1% FDR).*

*Supplementary Table 5: Comparison of pAla/pLeu/pGly Decoy FLR site counts searching PXD008355 (Arabidopsis data set) using different pipelines: TPP, PEAKS, MaxQuant and Mascot/ptmRS (PD) (fully tryptic, 1%FDR).*

|  | **Count at 1% FLR** | | | **Count at 5% FLR** | | | **Count at 10% FLR** | | |
| --- | --- | --- | --- | --- | --- | --- | --- | --- | --- |
|  | **pAla** | **pLeu** | **pGly** | **pAla** | **pLeu** | **pGly** | **pAla** | **pLeu** | **pGly** |
| **TPP** | 23104 | 17943 | 18872 | 40514 | 42157 | 35939 | 44556 | 45875 | 38885 |
| **PEAKS** | 4627 | 3633 | 1305 | 6679 | 5662 | 3778 | 13683 | 9725 | 4539 |
| **MaxQuant** | 9826 | 9086 | 6808 | 31104 | 18645 | 27503 | 33663 | 32436 | 29661 |
| **Mascot/ptmRS** | 23646 | 27294 | 20976 | 43840 | 43918 | 36414 | 57305 | 57591 | 54720 |

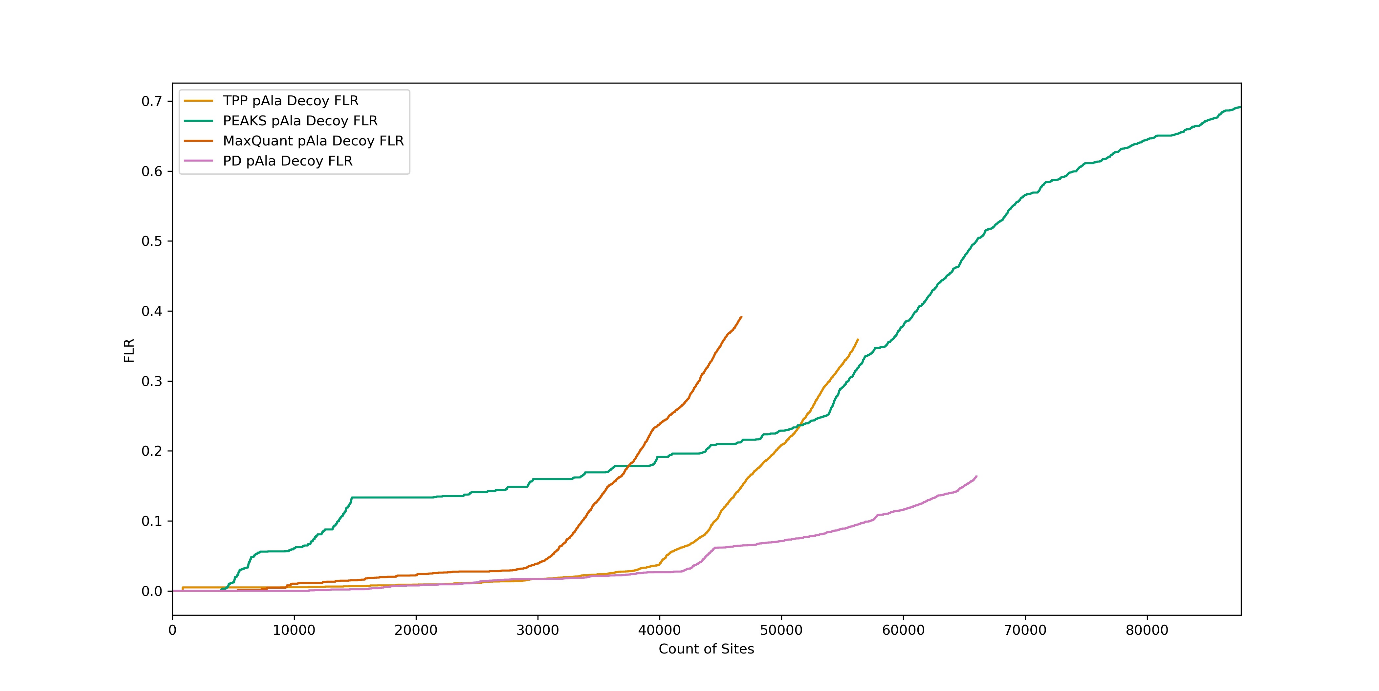

a)

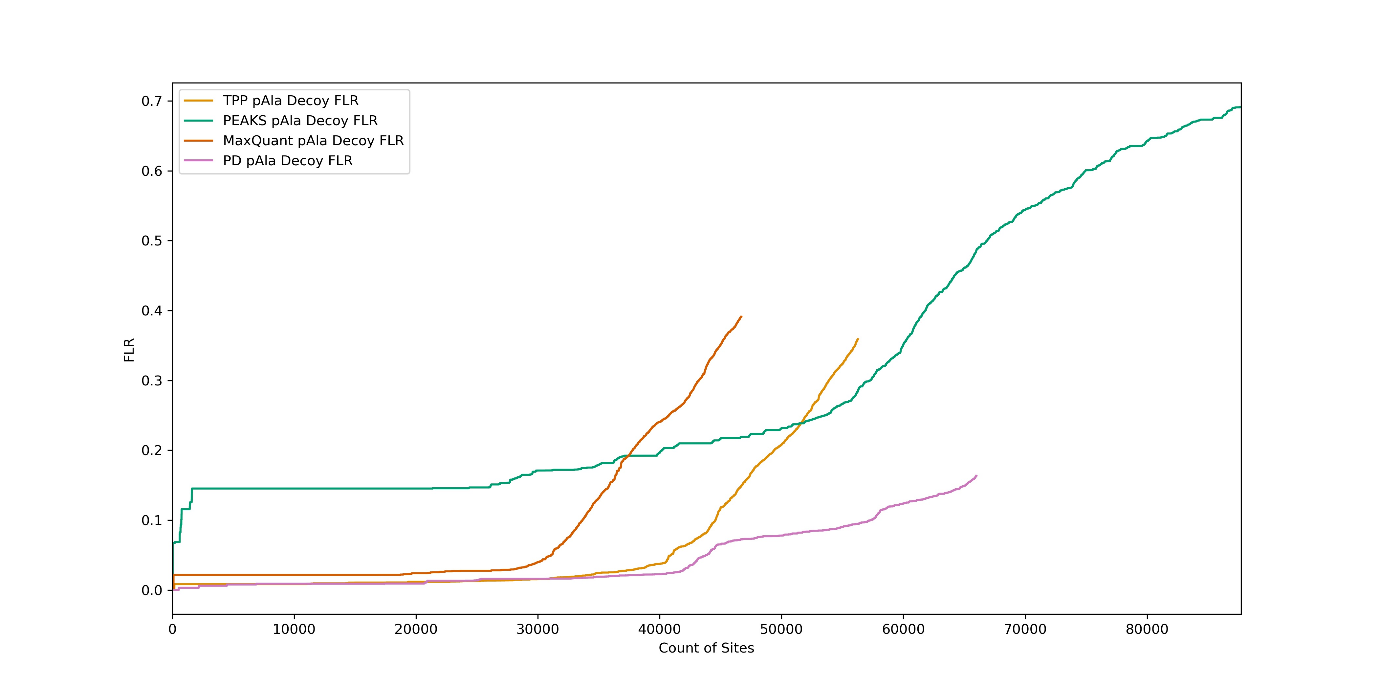

b)

*Supplementary Fig 7: Comparison of pAla Decoy FLR estimation searching PXD008355 (Arabidopsis data set) using different pipelines: TPP, PEAKS, MaxQuant, Mascot/ptmRS (PD). a) ordered by combined probability, b) ordered by PTM probability (Fully tryptic, 1 %FDR).*

*Supplementary Table 6: Counts of sites at pAla Decoy FLR for 1%, 5% and 10% using each pipeline, searching PXD008355 (Arabidopsis data set) (fully tryptic, 1% FDR).*

|  | **Ordered by combined probability** | | | **Ordered by PTM probability** | | |
| --- | --- | --- | --- | --- | --- | --- |
| **Pipeline** | **Count at 1% FLR** | **Count at 5% FLR** | **Count at 10% FLR** | **Count at 1% FLR** | **Count at 5% FLR** | **Count at 10% FLR** |
| **TPP** | 23104 | 40514 | 44556 | 1468 | 40921 | 44576 |
| **PEAKS** | 4627 | 6679 | 13683 | 23 | 23 | 734 |
| **MaxQuant** | 9826 | 31104 | 33663 | 91 | 31089 | 33667 |
| **Mascot/ptmRS** | 23646 | 43840 | 57305 | 20712 | 43881 | 57353 |

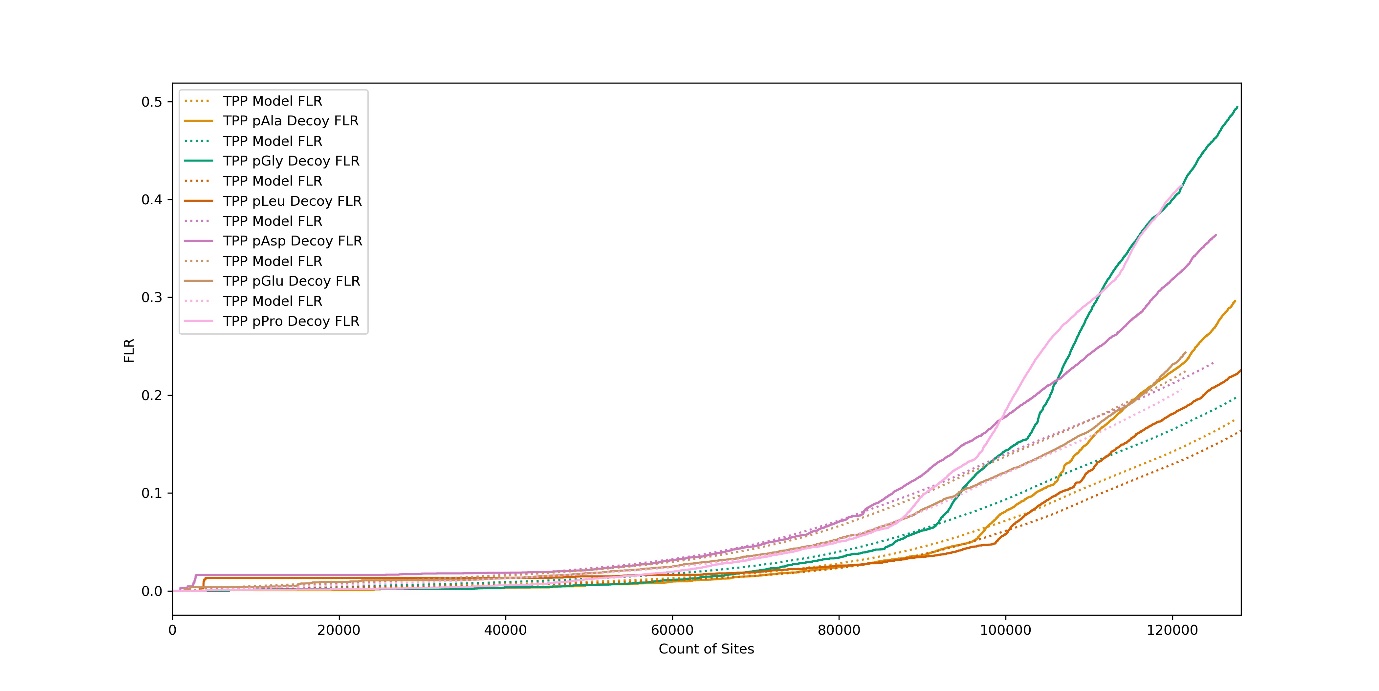

*Supplementary Fig 8: Comparison of pX Decoy FLR and Model FLR estimation searching PXD000612 (Human data set) for pAla, pLeu, pGly, pAsp, pGlu and pPro (Fully tryptic, 1 %FDR), using TPP.*

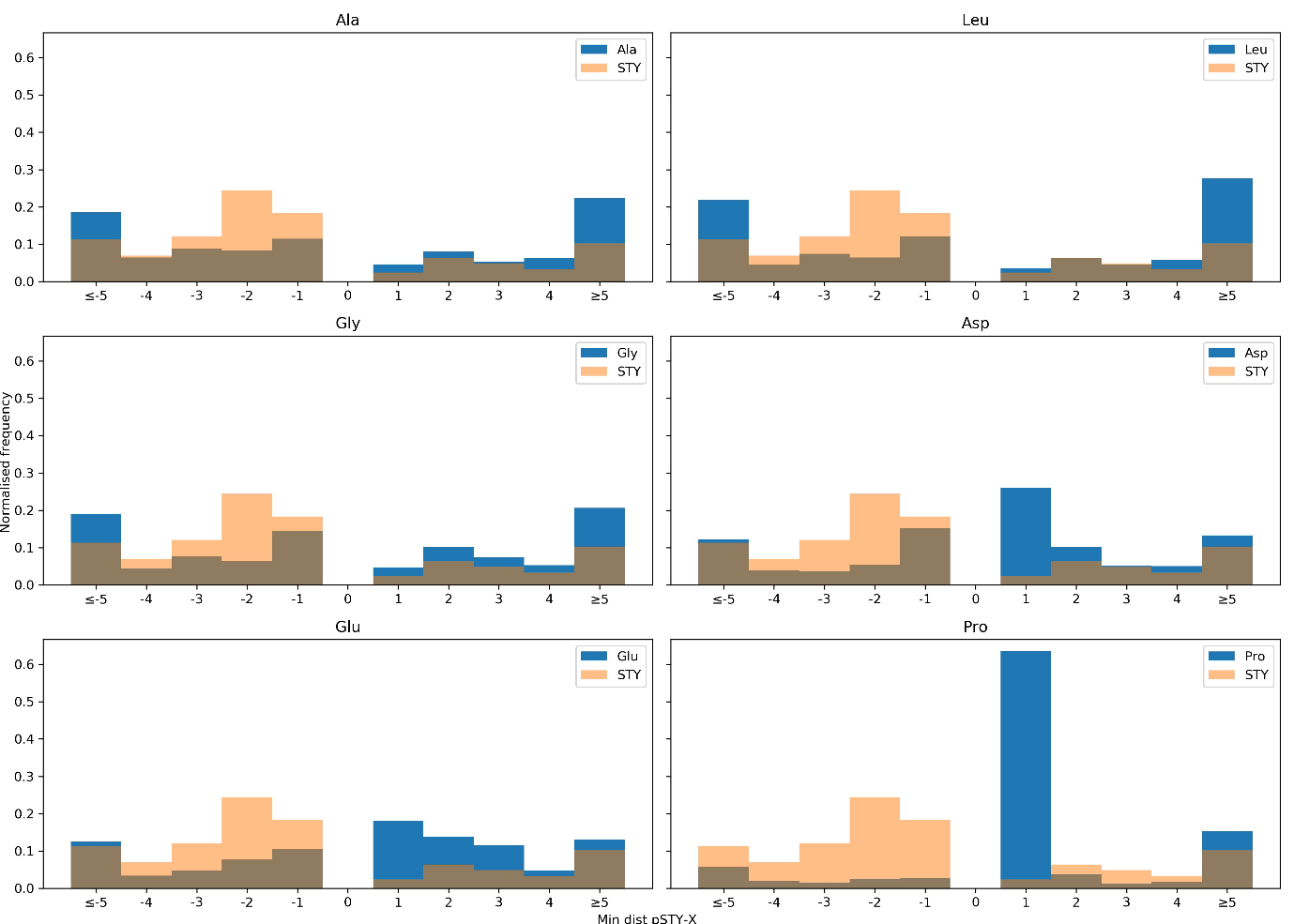

*Supplementary Figure 9: Comparison of minimum distance between phosphorylated S, T or Y and the nearest target amino acid (Ala, Leu, Gly, Asp, Glu and Pro), compared to the STY distribution, searching PXD000612 (Human data set).*

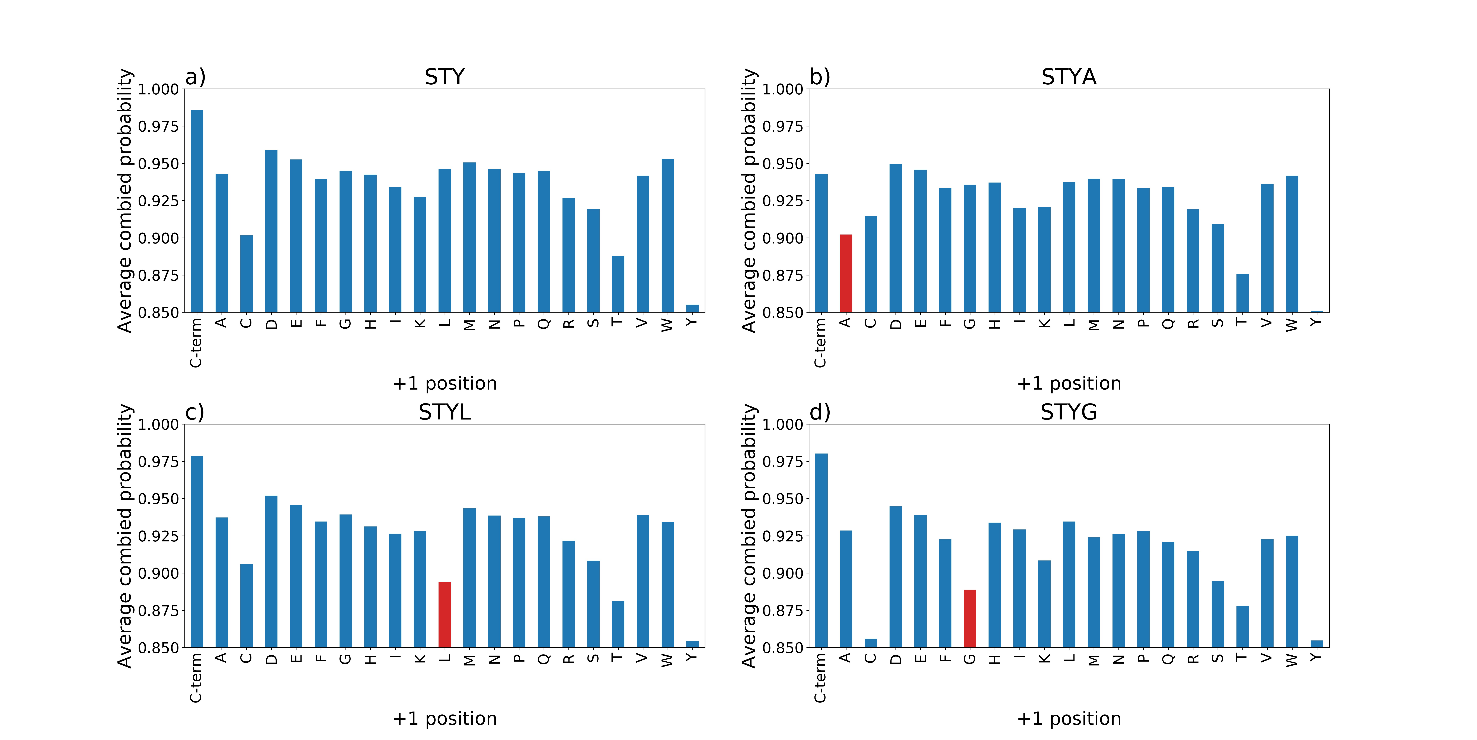

*Supplementary Figure 10: Comparison of averaged final site probabilities for all peptides (final probability ≥0.68), split by amino acid in the +1 positions for the PXD008355 (Arabidopsis data set) a) STY (no decoy), b) STY with Ala decoy, c) STY with Leu decoy and d) STY Gly decoy.*

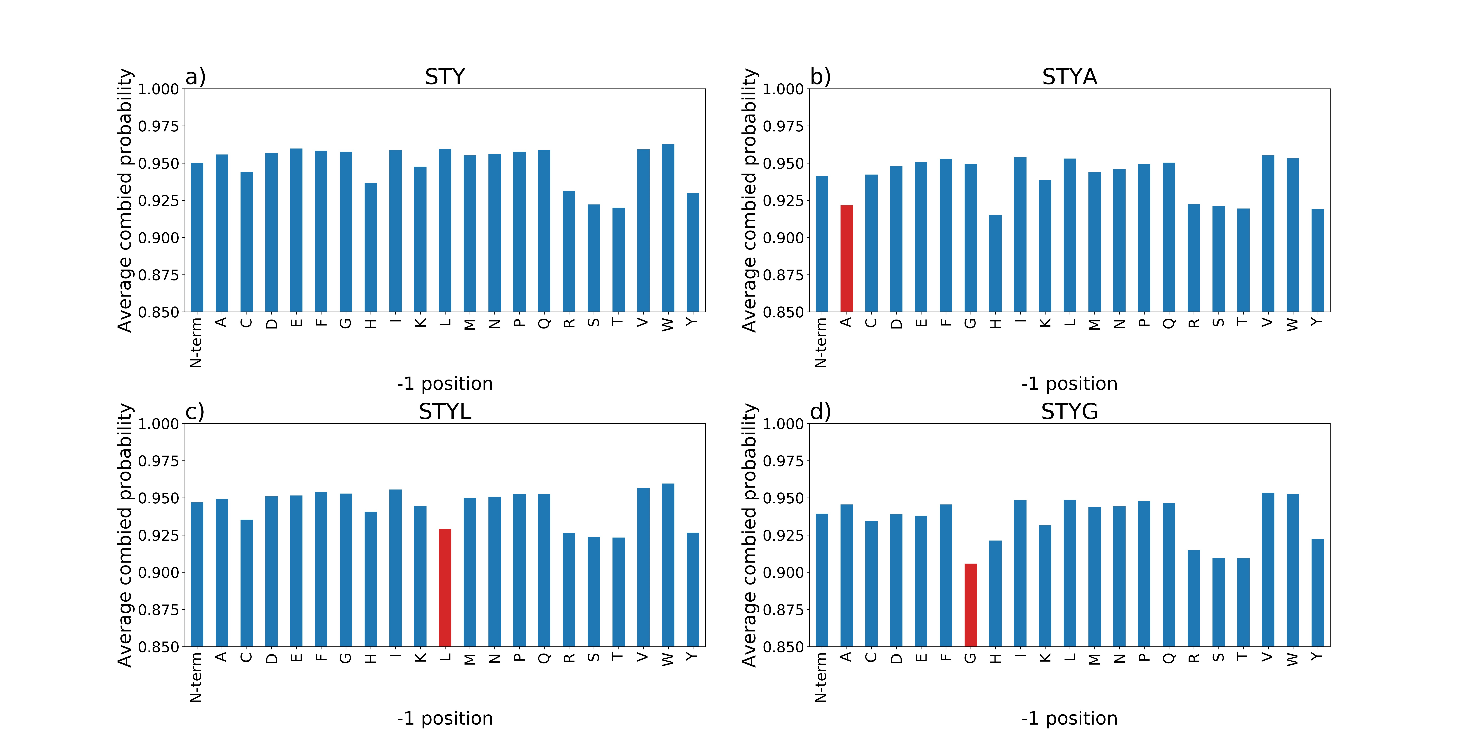

*Supplementary Figure 11: Comparison of averaged final site probabilities for all peptides final probability ≥0.68), split by amino acid in the -1 positions for the PXD000612 (Human data set) a) STY (no decoy), b) STY with Ala decoy, c) STY with Leu decoy and d) STY Gly decoy.*

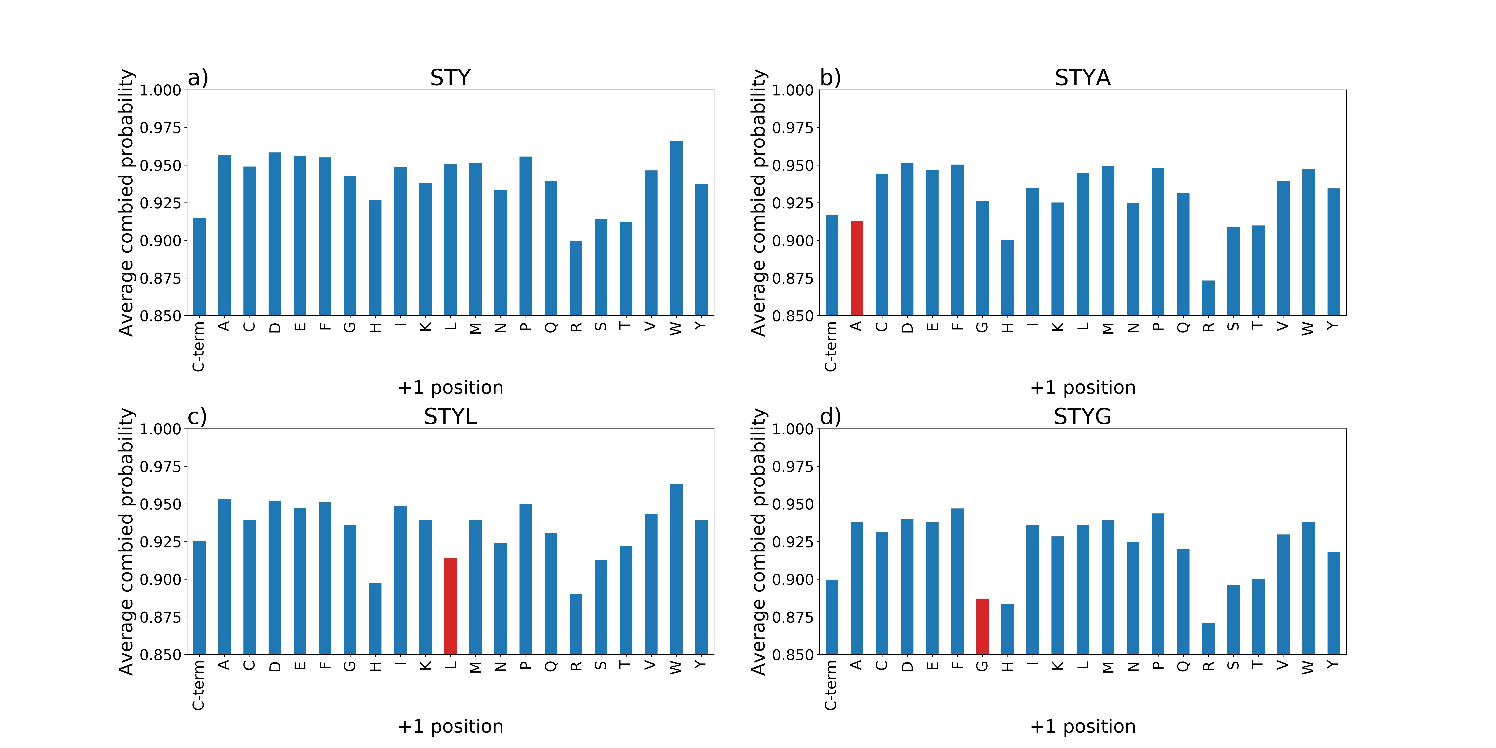

*Supplementary Figure 12: Comparison of averaged final site probabilities for all peptides (final probability ≥0.68), split by amino acid in the +1 positions for the PXD000612 (Human data set) a) STY (no decoy), b) STY with Ala decoy, c) STY with Leu decoy and d) STY Gly decoy.*
